# The Histone H3-H4 Tetramer is a Copper Reductase Enzyme

**DOI:** 10.1101/350652

**Authors:** Narsis Attar, Oscar A. Campos, Maria Vogelauer, Chen Cheng, Yong Xue, Stefan Schmollinger, Nathan V. Mallipeddi, Brandon A. Boone, Linda Yen, Sichen Yang, Shannon Zikovich, Jade Dardine, Michael F. Carey, Sabeeha S. Merchant, Siavash K. Kurdistani

**Affiliations:** Department of Biological Chemistry, David Geffen School of Medicine, University of California Los Angeles, Los Angeles, CA 90095, USA; Molecular Biology Institute, University of California Los Angeles, Los Angeles, CA 90095, USA; Institute for Genomics and Proteomics, Department of Chemistry and Biochemistry, University of California Los Angeles, Los Angeles, CA 90095, USA; Department of Molecular, Cell, and Developmental Biology, University of California Los Angeles, Los Angeles, CA 90095, USA; Eli and Edythe Broad Center of Regenerative Medicine and Stem Cell Research, David Geffen School of Medicine, University of California Los Angeles, Los Angeles, CA 90095, USA

## Abstract

Ancestral histones were present in organisms with small genomes, no nucleus, and little evidence for epigenetic regulation, suggesting histones may have additional older functions. We report that the histone H3-H4 tetramer is an enzyme that catalyzes the reduction of Cu^2+^ to Cu^1+^ when assembled *in vitro* from recombinant histones. Mutations of residues in the putative active site at the interface of the apposing H3 proteins alter the enzymatic activity and cellular processes such as Sod1 function or mitochondrial respiration that depend on availability of reduced copper. These effects are not due to altered gene expression or copper abundance but are consistent with decreased levels of cuprous ions. We propose that the H3-H4 tetramer is an oxidoreductase that provides biousable copper for cellular and mitochondrial chemistry. As the emergence of eukaryotes coincided with the Great Oxidation Event and decreased biousability of metals, the histone enzymatic function may have facilitated eukaryogenesis.

## Introduction

Eukaryotes owe their nucleosomal chromatin structure to an ancestral histone-containing archaeon that merged with the protomitochondria, forming the first eukaryotic common ancestor. Archaeal histone tetramers wrap ∼60 bp of DNA (Pereira et al., 1997), and have similar geometry and protein-DNA interactions (Mattiroli et al., 2017) as the eukaryotic H3-H4 tetramer. Unlike eukaryotic histones, archaeal histones typically lack extended N-terminal tails, are devoid of post-translational modifications, and occur in organisms with no known protein machinery for epigenetic regulation. Archaea also have significantly smaller genomes than eukaryotes, and do not contain nuclei. With no apparent capability for epigenetic regulation or little need for genome compaction in a confined nuclear space, the possibility arises that the conserved histone H3-H4 tetramer complex may have an additional older function in archaea, and therefore in the eukaryotes that emerged from them.

An overlooked feature of the nucleosome and the geochemical events surrounding eukaryogenesis hinted at what such a function might be. The region of the nucleosome where the two histone H3 proteins form a dimerization interface contains amino acid residues, including cysteine 110 (H3C110) and histidine 113 (H3H113), that are common in transition metal binding sites, particularly those that coordinate copper ions (Katz et al., 2003). These residues display some ability to coordinate transition metal ions *in vitro*, but it is unclear whether such interactions are functionally significant (Adamczyk et al., 2007; Bal et al., 1995; Saavedra, 1986). The evolutionary conservation of residues in this region is greater than expected given their contributions to the thermodynamic stability of the nucleosome (Ramachandran et al., 2011), leading to the suggestion that “a novel function” provided by the H3-H3’ interface is driving the unexpected conservation (Ramachandran et al., 2011). Whether the putative metal-binding capability of histone H3 is biologically relevant in eukaryotes is not known, but a clue comes from the geochemical history of Earth. The emergence of eukaryotes approximately coincided with the initial accumulation of molecular oxygen (Anbar, 2008). Its increasing abundance substantially altered the biousability of transition metals such as iron and copper in the oceans. These essential metals were also becoming more toxic in the presence of oxygen (Saito et al., 2003), presenting a formidable challenge for their usage. The hypothesis that early eukaryotes may have relied partly on histones for proper metal homeostasis has not been considered.

Copper is an essential element in eukaryotes that serves as a redox co-factor for numerous critical enzymes such as cytochrome *c* oxidase in the mitochondrial electron transport chain (ETC), and Cu, Zn-superoxide dismutases (e.g., Sod1) (Nevitt et al., 2012). Copper also functions as a signaling element to regulate specific biochemical pathways (Krishnamoorthy et al., 2016). In the active sites of redox enzymes, copper cycles between the cupric (Cu^2+^) and cuprous (Cu^1+^) forms, including partial oxidation states in between. However, it is the fully reduced cuprous form that is trafficked intracellularly, suggesting a need to maintain cellular cuprous ion pools for proper distribution and utilization of copper (Pufahl et al., 1997). How cells maintain a pool of cuprous ions is not fully understood. Here, we present biochemical, molecular, and genetic data that eukaryotic histone H3-H4 tetramers catalyze reduction of cupric to cuprous ions with the catalytic site likely forming at the H3-H3’ interface. Our data indicate that histones provide cuprous ions for utilization by various cellular and mitochondrial proteins. This function may have contributed to the emergence of eukaryotes, making the presence of histones in the ancestral archaeon not incidental (Sandman and Reeve, 1998), but rather essential for eukaryogenesis.

## Results

### Recombinant histone H3-H4 tetramer interacts with cupric ions

We first sought to determine whether copper ions interact with the residues at the putative metal-binding site of the H3-H3’ interface (Saavedra, 1986) (Figure 1A). We assembled and purified *Xenopus laevis* recombinant histone H3-H4 tetramers (Dyer et al., 2004), which are identical to human H3.2 and H4 histone proteins, respectively (Figure 1 – figure supplement 1A-C). We performed tetramer assembly and subsequent assays in solutions and glassware thoroughly depleted of contaminating metal ions, a critical precaution for *in vitro* studies (see Methods).

**Figure 1.**
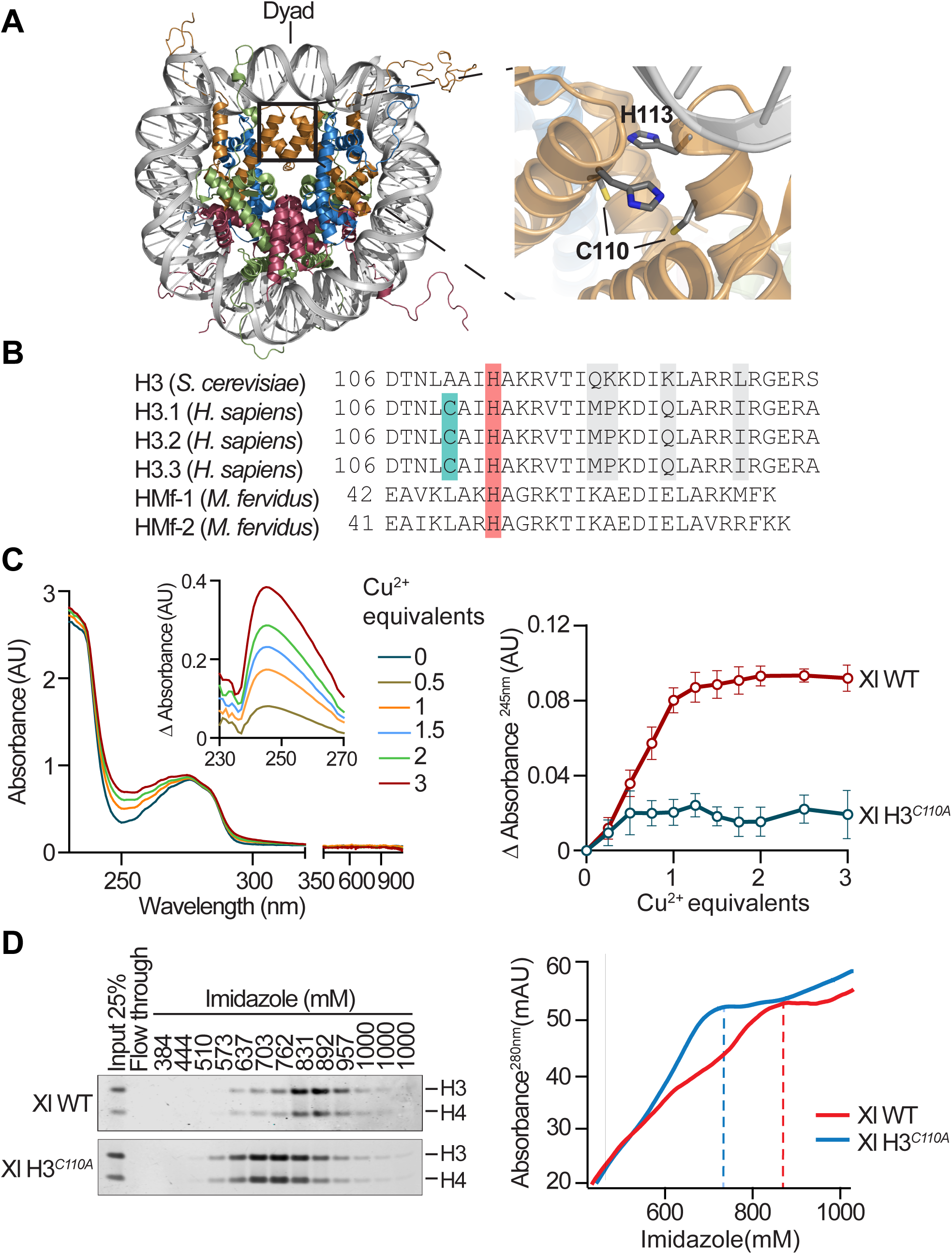
Recombinant histone H3-H4 tetramer interacts with cupric ions. **(A)** *Left*: *X. laevis* nucleosome core particle structure (PDB:1KX5) (Davey et al., 2002) viewed down the DNA superhelix axis. Histones H3, H4, H2A and H2B are shown in metallic orange, blue, green and red, respectively. The square box delineates the H3-H3’ interface. *Right*: Interface residues H113 and C110, one from each of the histone H3 proteins, are highlighted. **(B)** Alignment of the C-terminal region of *S. cerevisiae* histone H3, human non-centromeric histone H3 variants, and M. *fervidus* HMf-1 and HMf-2. **(C)** *Left panel*: UV-vis absorbance spectrum of the *X. laevis* H3-H4 tetramer incubated with increasing molar equivalents of Cu^2+^. Inset: Differential UV absorbance compared to the absorbance of the tetramer alone, and without subtraction of buffer absorbance. *Right panel*: Buffer-corrected differential UV absorbance at 245nm of the indicated *X. laevis* tetramers incubated with increasing molar equivalents of Cu^2+^. **(D)** Elution of the indicated *X. laevis* tetramers from a Cu^2+^-IMAC column. *Left panel*: Image of coomassie stained PAGE of input (25%), flow-through, and fractions eluted at indicated concentrations of imidazole. *Right panel*: corresponding FPLC elution profile over a gradient of increasing imidazole concentrations. Peaks of tetramers appear over background absorbance of increasing imidazole concentrations (i.e., lines trending up).

We focused first on the role of the pair of cysteine residues at position 110 (H3C110), one from each apposing H3 protein (Adamczyk et al., 2007; Saavedra, 1986), in copper ion binding. H3C110 is present in the canonical histone H3 of 146 out of 166 eukaryotes that span the major kingdoms (Macadangdang et al., 2014), including *X. laevis*, and *H. sapiens.* It is the sole cysteine residue present in most H3s or in H4 and is absent only in a few fungi of the genus Saccharomyces, including *S. cerevisiae* H3 (Figure 1B). Cysteine-copper ion interactions generally result in characteristic absorption spectra in the UV-visible range that are dependent on the coordination environment (Solomon et al., 1993). Incubation of the *X. laevis* wildtype (Xl WT) tetramer with increasing molar equivalents of Cu^2+^ resulted in an increasing UV absorbance band with a peak at 245 nm that plateaued at a tetramer to cupric ion ratio of 1 (Figure 1C, left panel). Critically, mutation of C110 to alanine (*H3C110A*), which did not affect tetramer assembly (Figure 1 – figure supplement 1B and C), abolished this copper-dependent absorbance change (Figure 1C, right panel), consistent with a direct interaction between C110 and a copper ion. Furthermore, when assayed by Immobilized Metal Affinity Chromatography (IMAC) using a Cu^2+^-loaded resin, the Xl WT tetramer exhibited substantially more retention than the H3*^C110A^* tetramer, corroborating the role of H3C110 in tetramer-copper ion interaction (Figure 1D).

### The histone H3-H4 tetramer is a cupric ion reductase

We next hypothesized that the H3-H4 tetramer may not only bind copper but function as an oxidoreductase, catalyzing the reduction of Cu^2+^ to Cu^1+^ in the presence of a source of electrons. We developed a colorimetric assay to measure the production of Cu^1+^ by utilizing copper chelators neocuproine (NC) or bicinchoninic acid (BCA) to detect Cu^1+^ quantitatively (Figure 2A, Figure 2 – figure supplement 1, and Figure 2 – figure supplement 2A and B). Formation of the NC_2_-Cu^1+^ or the BCA_2_-Cu^1+^ complex result in a change in absorbance at 448 nm or 562 nm, respectively. The assay also contained reduced forms of tris(2-carboxyethyl)phosphine (TCEP), nicotinamide adenine dinucleotide (NADH), or its phosphate form (NADPH) as electron donors. Reactions were initiated by addition of Cu^2+^ substrate in the form of a CuCl_2_-Tricine, CuCl_2_-N-(2-Acetamido)iminodiacetic acid (ADA), or CuCl_2_-Nitrilotriacetic acid (NTA) solution, upon which spontaneous Cu^1+^ production occurred at a slow rate in control buffers (Figure 2B). In contrast, the rate of Cu^1+^ production was substantially increased when reactions were performed in the presence of the Xl WT tetramer (Figure 2B). Production of Cu^1+^ plateaued at later time points due to near full consumption of the electron donor with ∼75% of total Cu^1+^ produced in the first 30 seconds (Figure 2B). Importantly, the rate of Cu^1+^ production was decreased to near-background levels upon heat-inactivation of the Xl WT tetramer (Figure 2B). The unassembled *X. laevis* histone H3 on its own or another histone, the yeast H2A, did not enhance cupric ion reduction (Figure 2 – figure supplement 2C). Digestion of the H3-H4 tetramer with Proteinase K prior to the assay abolished cupric reductase activity (Figure 2C), confirming the importance of the structural conformation of the tetramer for copper reduction. No significant production of Cu^1+^ occurred in the absence of TCEP and increasing rates of production occurred with increasing amounts of either TCEP (Figure 2D) or Cu^2+^ (Figure 2E), eventually approaching a maximum rate of reduction which is consistent with protein-based catalysis (Figure 2 – figure supplement 2D and E). The tetramer also enhanced the rate of cupric ion reduction with NADPH, albeit in a different reaction condition that was necessary to decrease un-catalyzed copper reduction by NADPH (Figure 2F).

**Figure 2.**
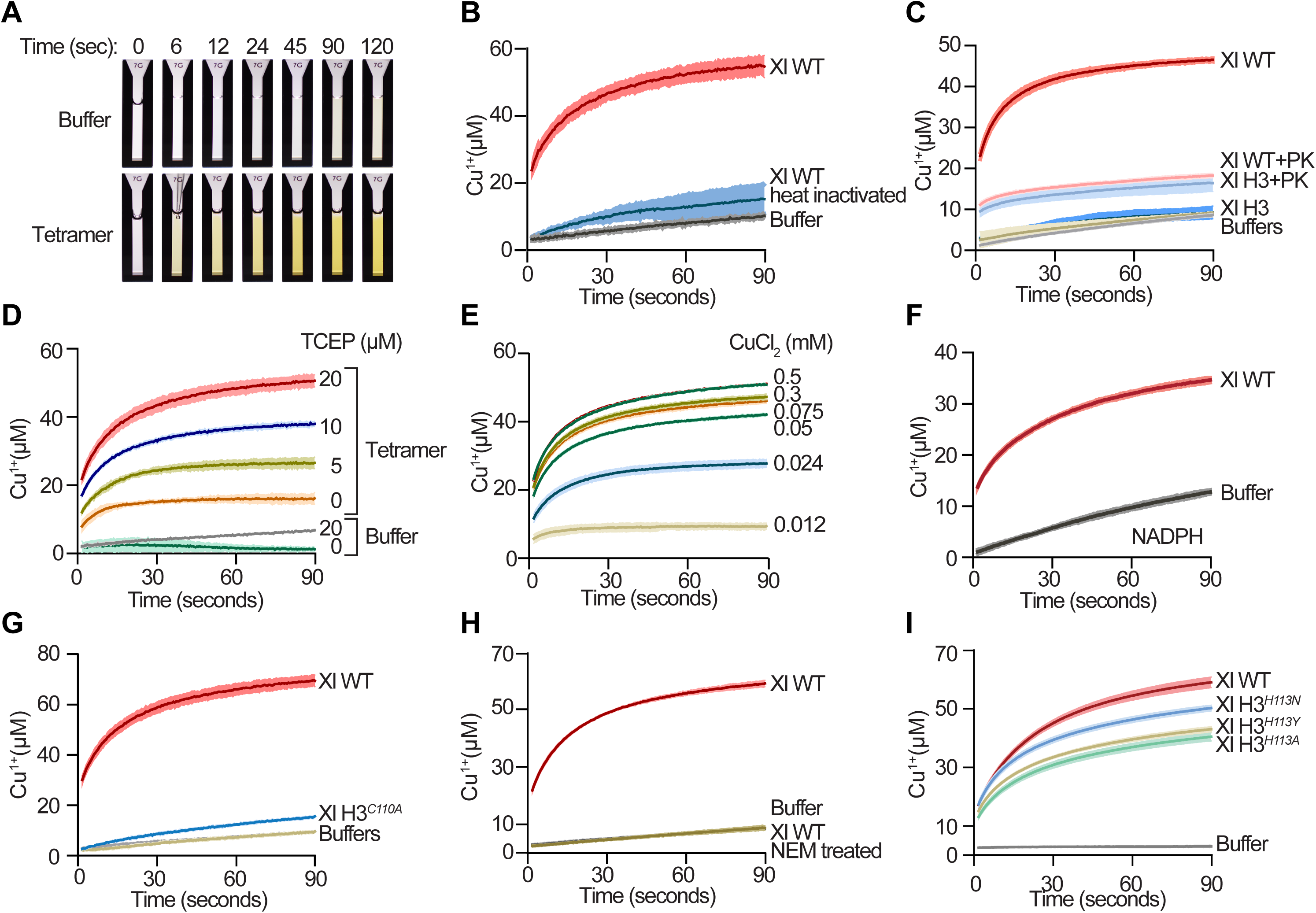
The *Xenopus laevis* H3-H4 tetramer is a copper reductase. **(A)** Photographic representation of *in vitro* copper reductase assay at indicated times. Control buffer or *X. laevis* tetramer were reacted with 30 µM TCEP and 1 mM CuCl_2_ in 10 mM Tricine pH 7.5 in presence of neocuproine (NC) Cu^1+^ chelator. The yellow color is due to NC_2_-Cu^1+^ complex formation. **(B)** Progress curves of copper reduction by 1 µM of *X. laevis* H3-H4 tetramer, heat-inactivated tetramer, or buffer in presence of 30 µM TCEP and 1 mM CuCl_2_ in 10 mM Tricine. Lines and shading represent the mean ± SD of 3-5 assays. **(C)** Same as (B) using 1 µM of *X. laevis* tetramer and equimolar monomeric *X. laevis* H3 before and after digestion with Proteinase K (PK). **(D)** Same as (B) with indicated TCEP concentrations. **(E)** Same as (B) with indicated CuCl_2_ concentrations. **(F)** Progress curves of copper reduction by 1 µM of *X. laevis* tetramer using 20 µM NADPH as the electron donor with 0.5 mM CuCl_2_ in 7.5 mM Tricine pH 7.5. Lines and shading represent the mean ± SD of 3-5 assays. **(G)** Same as (B) with the indicated *X. laevis* tetramers. **(H)** Same as (B) with and without NEM treatment of the tetramer. **(I)** Progress curves of copper reduction by 5 µM of the indicated *X. laevis* tetramers in presence of 100 µM TCEP and 0.5 mM CuCl_2_ in 0.5 mM ADA pH 8.

The *C110A* mutation in Xl H3 substantially reduced the rate of cupric ion reduction (Figure 2G). Treatment of the Xl WT tetramer with *N*-ethylmaleimide (NEM) to irreversibly inhibit H3C110 reactivity abolished Cu^1+^ production (Figure 2H), corroborating the importance of H3C110 in cupric ion reduction. Given the general ability of thiols to reduce cupric ions *in vitro*(Pecci et al., 1997), we considered whether any set of protein or small molecule thiol groups may enhance the rate of cupric ion reduction by TCEP, as opposed to merely donating electrons directly to Cu^2+^. Reactions carried out in the presence of RNase A, which contains eight redox reactive cysteines capable of forming four disulfide bonds (Klink et al., 2000), at the same concentration as the tetramer did not enhance the rate of copper reduction (Figure 2 – figure supplement 2F). Similarly, 1 µM of DTT directly reduced an equimolar 2 µM of Cu^2+^ but did not substantially enhance the rate of cupric ion reduction by TCEP (Figure 2 – figure supplement 2F), emphasizing the unique role of H3C110 in the structural context of the H3-H4 tetramer for enzymatic activity.

The pair of histidine residues at position 113 (H3H113) at the H3-H3’ interface have also been proposed to participate in metal coordination (Adamczyk et al., 2007; Saavedra, 1986). This highly conserved residue is present in the canonical histone H3 of all 166 eukaryotes that span the major kingdoms (Macadangdang et al., 2014), as well as in the structurally-equivalent position of most (29 out of 33) archaeal histones spanning the major phyla (Henneman et al., 2018) (Figure 1A and B). Thus, we asked whether the H3H113 residue also enables the H3-H4 tetramer to enhance the rate of cupric ion reduction. Mutation of H3H113 to alanine (*H3H113A*) or to asparagine or tyrosine (*H3H113N* or *H3H113Y*), which have been found in certain cancers (Forbes et al., 2015), had little effect on the rate of reduction with Cu^2+^-Tricine as the substrate (data not shown) but diminished cupric ion reduction in reactions with Cu^2+^-ADA (Figure 2I and Figure 2 – figure supplement 2G) or Cu^2+^-NTA as Cu^2+^ substrates (Figure 2 – figure supplement 2H and I). The differing effects of ADA, NTA, and Tricine on copper reduction rates are likely due in part to their differing Cu^2+^ coordination ability and/or geometry. Altogether, our data reveal that the H3-H4 tetramer uses electron donors as cofactors to catalyze Cu^2+^ reduction and is thus an oxidoreductase enzyme.

The histone H3 of *S. cerevisiae* naturally lacks the C110 residue found in most other eukaryotic H3s, and instead contains an alanine at this position (Figure 1B). It does, however, contain the H3H113 residue at the H3-H3’ interface. As expected, recombinant yeast H3-H4 tetramer did not absorb UV energy in the 245 nm range when incubated with Cu^2+^ (see below), as this was associated with the H3C110-copper ion interaction (Figure 1C). However, a broad peak centered at 680 nm was observed with increasing molar equivalents of Cu^2+^, which required high concentrations of yeast tetramer, consistent with weakly-absorbing d-d transitions that are typical of coordinated Cu^2+^ ions (Mesu et al., 2006) (Figure 3A, left panel). Notably, this absorbance band only occurred when the yeast tetramer was assembled and incubated with Cu^2+^ ions in the presence of 2 M NaCl, but not at lower ionic strengths (data not shown), which may interfere with the structural conformation of the yeast tetramer (Lee et al., 1982). The *H113A* mutation in the yeast H3, which did not affect tetramer assembly (Figure 3 – figure supplement 1A and B), strongly reduced copper-dependent absorbance change at 680 nm (Figure 3A, right panel), consistent with the role of the histidine in copper ion interaction independently of the H3C110 residue. The yeast H3-H4 tetramer also exhibited substantial retention on the Cu^2+^ IMAC column, which was reduced in tetramers harboring the *H3H113A, N,* or *Y* mutations (Figure 3B and Figure 3 – figure supplement 1A-D).

**Figure 3.**
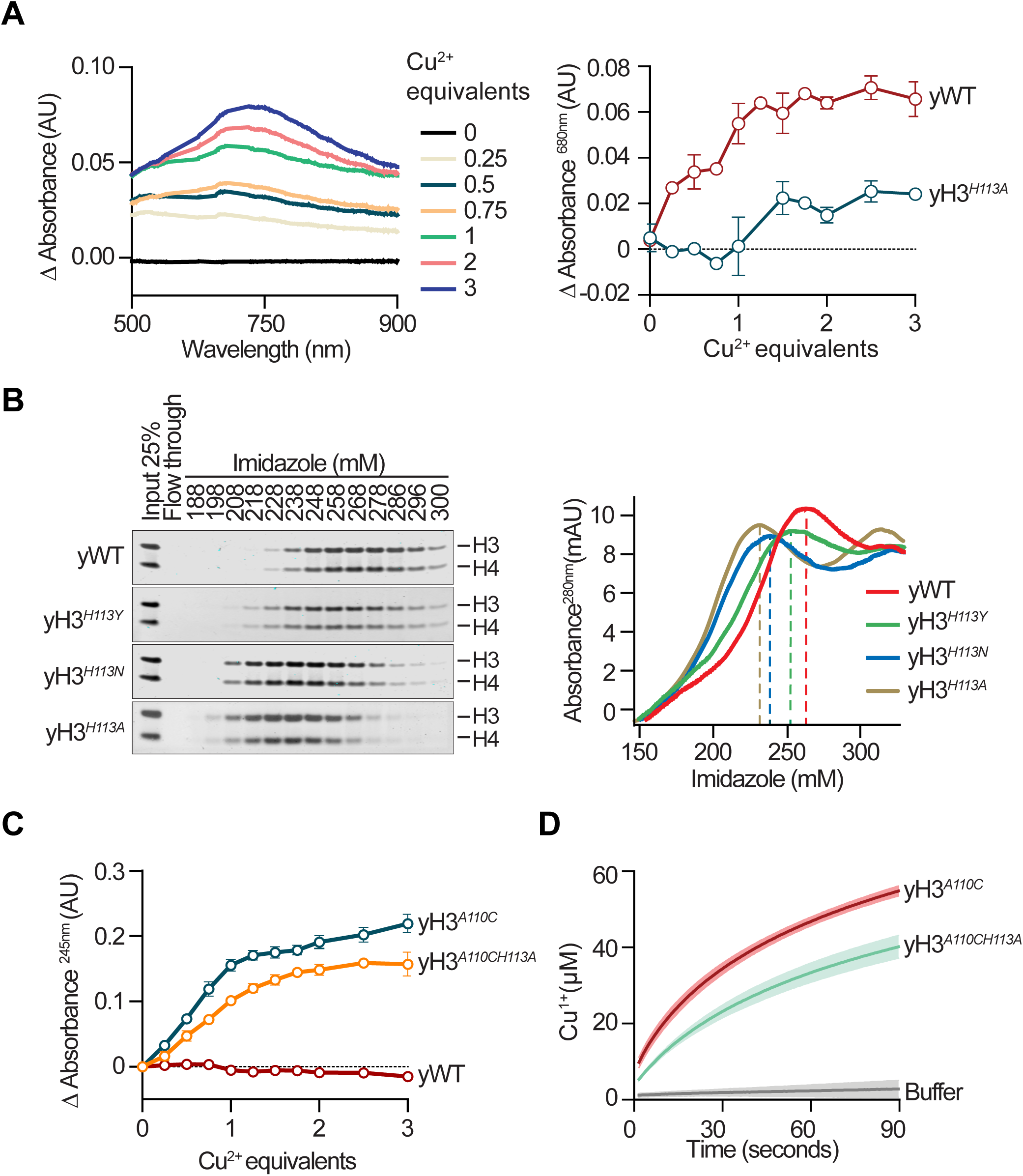
The *Saccharomyces cerevisiae* H3-H4 tetramer potentially is a copper reductase. **(A)** *Left panel*: Differential absorbance of visible light between 500 and 900 nm compared to the absorbance of the yeast tetramer alone, and without subtraction of buffer absorbance. *Right panel*: Buffer-corrected differential red light absorbance at 680 nm of the indicated yeast tetramers incubated with increasing molar equivalents of Cu^2+^. **(B)** Elution of the indicated *S. cerevisiae* tetramers from a Cu^2+^-IMAC column. *Left panel*: Image of coomassie stained PAGE of input (25%), flow-through, and fractions eluted at indicated concentrations of imidazole. *Right panel*: corresponding FPLC elution profile over a gradient of increasing imidazole concentrations. Peaks of tetramers appear over background absorbance of increasing imidazole concentrations (i.e., lines trending up). **(C)** Buffer-corrected differential UV absorbance at 245 nm of the indicated *S. cerevisiae* tetramers incubated with increasing molar equivalents of Cu^2+^. **(D)** Progress curves of copper reduction by 5 µM of the indicated *S. cerevisiae* tetramers in presence of 100 µM TCEP and 0.5 mM CuCl_2_ in 0.5 mM ADA pH 8.

Mutation of yeast H3A110 to cysteine (yH3*^A110C^* tetramer) (Figure 3 – figure supplement 1E and F) elicited a copper-dependent absorbance band centered at 245 nm, nearly identical to that observed with the *X. laevis* tetramer (Figure 3C), corroborating the ability of the yeast tetramer to interact with copper ions at the H3-H3’ interface. Furthermore, the *H3H113A* mutation in the context of the yeast H3*^A110C^* tetramer (yH3*^A110CH113A^* tetramer) (Figure 3 – figure supplement 1E and G) resulted in diminished copper-dependent absorbance at 245 nm (Figure 3C), indicating that H3H113 affects the nature of the cysteine-copper ion interaction. Like the *X. laevis* tetramer, the yeast H3*^A110C^*-H4 tetramer displayed cupric reductase activity, which was diminished by the *H3H113A* mutation (Figure 3D). In the absence of the H3C110 residue, the yeast tetramer did not display robust cupric reductase activity in standard salinity (∼100-150 mM NaCl) likely due to deficient Cu^2+^ binding, unsatisfactory yeast tetramer conformation and/or inadequate reconstitution of the cellular environment in our assay conditions. We also could not perform copper reduction assays in high (2 M) NaCl concentrations, at which the yeast tetramer interacts with Cu^2+^ (Figure 3A), due to salt interference with other assay components. Nonetheless, our findings suggest that the yeast H3-H4 tetramer can display cupric reductase activity, and that the H3H113 residue participates in this function.

*(Note to reviewers: We have attempted to perform the curpric reductase assay with nucleosomes. However, the combination of Cu^2+^, the copper chelator, and DNA form an insoluble complex, precluding us from performing enzyme assays. We are working on this problem but may or may not make a breakthrough in a reasonable timeframe.)*

### Mutation of histone H3 histidine 113 results in loss of function in yeast

We next investigated whether mutation of the H3H113 residue *in vivo* would produce cellular effects consistent with the ability of histone H3 to produce Cu^1+^ ions. Mutation of H3H113 to alanine was lethal in *S. cerevisiae* (Huang et al., 2009) (Figure 4A). However, introduction of either *H3H113N* or *H3H113Y* mutations in the two chromosomal copies of the histone H3 gene generated viable yeast strains *H3^H113N^* and *H3^H113Y^*, albeit with slower growth rates compared to WT (Figure 4B). Although replacement of H3H113 with any of three residues resulted in loss of copper binding by the H3-H4 tetramers *in vitro*, we sought to rule out the possibility that the slow growth phenotypes were due to gain of a toxic function *in vivo*. Yeast strains in which the *H3H113N* and *H3H113Y* mutant histones were expressed from only one locus, with the other locus deleted, were severely sick compared to strains expressing both histone copies (Figure 4C), confirming that substitution of H3H113 with specific amino acids generates loss-of-function alleles.

**Figure 4.**
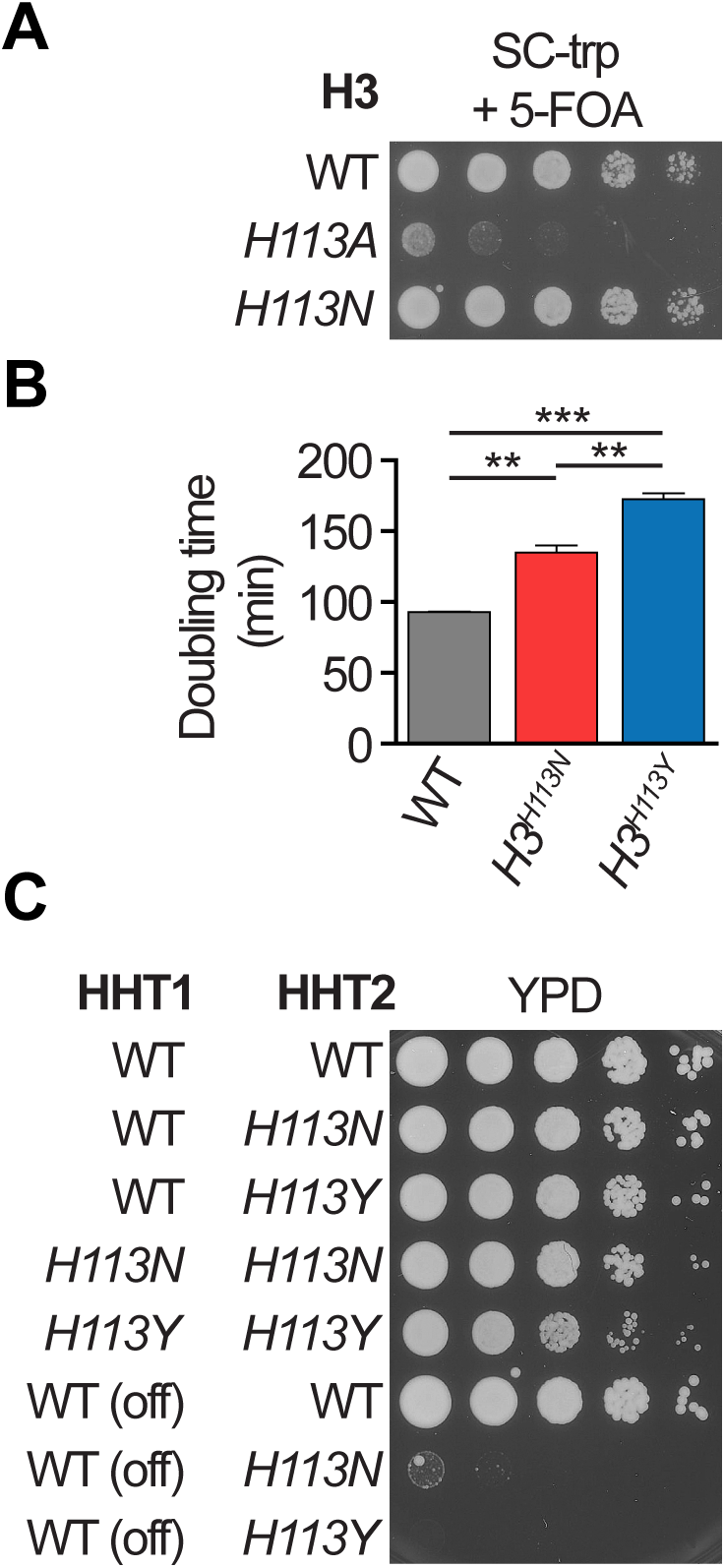
Mutation of histone H3 histidine 113 in *S. cerevisiae* is a loss of function. **(A)** Plasmid shuffle assay with strains harboring WT H3 on a URA3 plasmid and the indicated H3 gene on a TRP1 plasmid. 5-Fluoro-orotic acid (5-FOA) is lethal to cells bearing the URA3 plasmid, and therefore selects for cells that maintain only the indicated H3 plasmid. **(B)** Average doubling times from steady-state growth in liquid Synthetic Complete (SC) medium. Bar graphs show mean ± standard deviation (SD) from 3-9 replicate cultures. **(C)** Spot test assay in fermentative medium (YPD). Note that the *HHT1* gene in the bottom three strains was placed under the control of the *GAL1* promoter, which is repressed in YPD. Since one copy of *H3H113N* or *H3H113Y* (rows 7 and 8) causes a more severe phenotype than two copies (*H3^H113N^* and *H3^H113Y^*, rows 4 and 5), this suggests that they are likely loss of function mutations.

### H3H113 mutations lead to de-repression of Mac1 target genes

We next examined whether the *H3H113N* or *H3H113Y* mutations result in decreased cuprous ion availability. We first investigated the activity of the transcription factor Mac1 because it is directly inhibited by cuprous ions in the nucleus (Graden and Winge, 1997). Global gene expression analysis by mRNA-seq revealed largely similar profiles in WT and *H3^H113N^* strains in rich fermentative medium (YPD and SC) (Figure 5 – figure supplement 1). However, Mac1 target genes displayed increased expression in *H3^H113N^* compared to WT (Figure 5A), which was corroborated in the *H3^H113Y^* strain via gene-specific RT-qPCR analysis (Figure 5B). Deletion of *CTR1*, the main copper importer in yeast (Dancis et al., 1994), expectedly resulted in upregulation of Mac1 target genes (Figure 5A). However, Mac1 target genes showed greater upregulation as well as incomplete repression in response to exogenous copper in *H3^H113N^ctr1*Δ compared to *ctr1*Δ (Figure 5C). Importantly, although *ctr1*Δ substantially reduced total intracellular levels of copper, they were unchanged in strains with the *H3H113N* mutation compared to strains with WT H3 (Figure 5D). This finding suggests that the effect of H3H113 mutations on Mac1 activity is not due to altering copper abundance but is consistent with decreased availability of cuprous ions in the nucleus.

**Figure 5.**
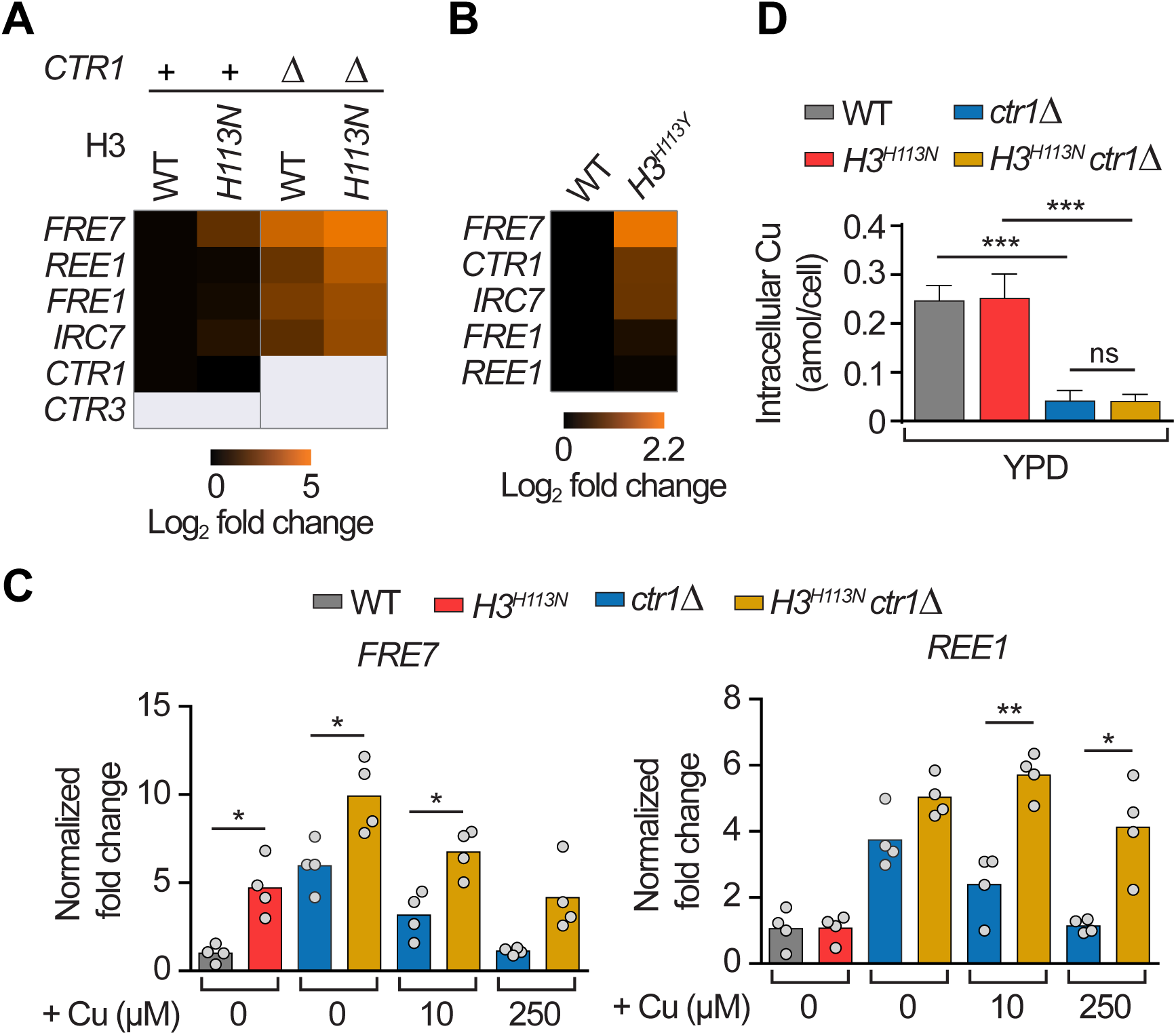
Mutation of H3H113 decreases the copper-dependent repression of Mac1 target genes. **(A)** Heat map of average fold changes of Mac1 target gene (Gross et al., 2000) expression, relative to WT for each gene and normalized to ACT1 fold changes, based on mRNA-seq analyses from two independent experiments. Genes have been ordered vertically based on the fold increase in the *H3^H113N^ctr1*Δ strain (4th column). Note that *CTR3* is a Mac1 target gene but is not expressed in this strain background due to a genetic disruption by a Ty2 transposon. **(B)** Same as (A) except showing fold changes in expression levels based on RT-qPCR analyses. The data are averages from four independent experiments in fermentative medium (YPD). Genes have been ordered vertically based on the fold increase in the *H3^H113Y^* strain. **(C)** Intracellular copper content measured by ICP-MS for exponentially growing strains in YPD. Data are presented as mean ± SD from 3-6 replicate cultures. **(D)** Fold changes in expression levels for two Mac1 target genes, normalized to ACT1, based on RT-qPCR analyses. Mac1 activation was induced by deletion of *CTR1* and repressed by 1 hr treatment with CuSO_4_ in YPD as indicated. Bars are the sample means and each dot is an independent experiment (n = 4). *P≤0.05, **P≤0.01, ***P<0.001.

### The H3H113 residue is required for efficient use of copper for mitochondrial respiration

We next tested whether mutation of H3H113 impacts mitochondrial respiration, which depends on copper ions for cytochrome *c* oxidase function. Although *H3^H113N^* did not display a defect in mitochondrial respiration, the more defective *H3^H113Y^* strain displayed a significant loss of cellular O_2_ consumption in respiratory media where glucose was replaced with the non-fermentable ethanol and glycerol (YPEG) (Figure 6A), consistent with its greater defect in Mac1 repression (Figure 5B). This defect was exacerbated in copper depleted medium, which was achieved by addition of the copper chelator bathocuproinedisulfonic acid (BCS), and was recovered by excess exogenous copper (Figure 6B), suggesting that copper utilization is perturbed when H3H113 is mutated.

**Figure 6.**
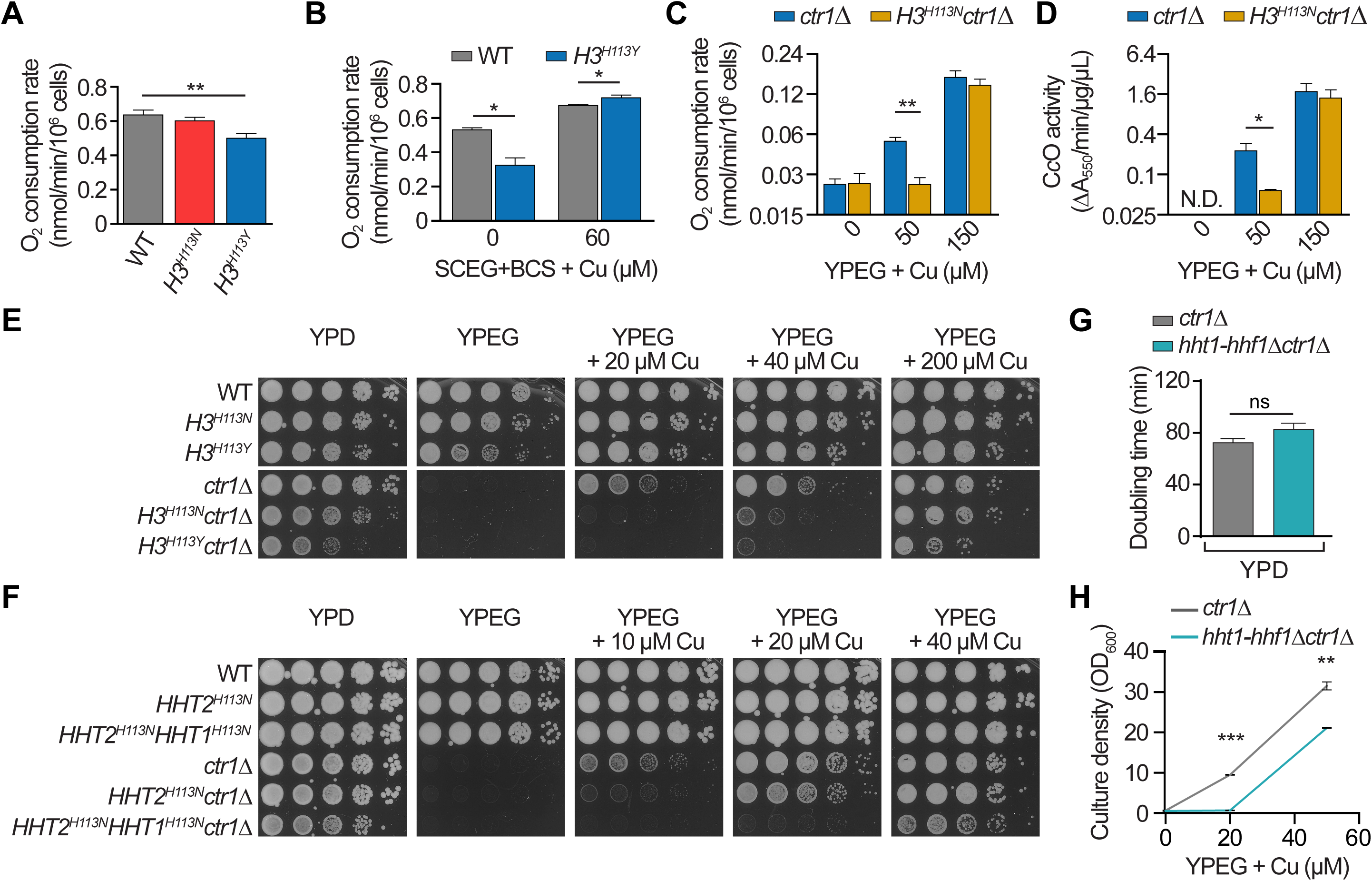
The H3H113 residue is required for efficient use of copper for mitochondrial respiration. **(A)** Oxygen consumption assays of cells incubated for 18 hrs in liquid YPEG. Baseline copper concentration in YPEG is ∼1 µM. Bars show means ± SD from three independent experiments done concurrently with the same batch of media and scaled to relative mitochondrial DNA contents (data not shown). **(B)** Same as (A) but with cells incubated in SCEG + 60 µM BCS. Baseline copper concentration in SCEG is ∼1 µM. **(C)** Oxygen consumption assays of cells incubated for 4 hrs in liquid YPEG with increasing amounts of CuSO_4_. Bars show means in log_2_ scale ± SD from three independent experiments. **(D)** Cytochrome *c* oxidase assays of cells incubated for 4 hrs in liquid YPEG with increasing amounts of CuSO_4_. Bars show means in log_2_ scale ± SD from three independent experiments. N.D.: not detectable. **(E)** Spot test assays in fermentative (YPD) or respiratory media (YPEG) with the indicated amounts of CuSO_4_. **(F)** Same as (E). **(G)** Average doubling times ± SD from steady-state growth in liquid YPD from three replicate cultures. **(H)** Average OD_600_ ± SD for growth in liquid YPEG with increasing amounts of CuSO_4_ after 36 hrs from three replicate cultures. *P≤0.05, **P≤0.01, ***P<0.001.

To relate defects in mitochondrial respiration to disruption of copper utilization, we assessed the ability of cells to increase respiratory function as they were shifted from a copper depleted state to a copper replete state. As expected, copper depletion via *CTR1* deletion resulted in a low rate of O_2_ consumption which was gradually increased by addition of copper exogenously (Figure 6C). However, substantially more exogenous copper was required to rescue *ctr1*Δ in the context of the *H3H113N* mutant histone (Figure 6C). The O_2_ consumption measured in our assays was almost entirely abolished by treatment with antimycin A (Figure 6 – figure supplement 1A), an inhibitor of cytochrome *c* reductase, reaffirming the specific impact of the *H3H113N* mutation on mitochondrial respiration. *H3^H113N^ctr1*Δ also displayed reduced recovery of copper-dependent cytochrome *c* oxidase activity compared to *ctr1*Δ (Figure 6D). The deficiencies in the context of *H3H113N* were apparent at early times after inoculation in liquid medium before any measurable growth had occurred, suggesting that they were not merely consequences of diminished growth rate. *CTR1* deletion abolished respiratory growth on YPEG, which was partially rescued by exogenous copper. Substantially more exogenous copper was required to rescue growth of *ctr1*Δ in the context of the *H3H113N* or *H3H113Y* mutant histones (Figure 6E and Figure 6 – figure supplement 1B). Combination of *H3H113N* with deletion of *MAC1*, which is required for *CTR1* expression (Graden and Winge, 1997), also increased the amount of exogenous copper required for growth on YPEG compared to *mac1*Δ alone (Figure 6 – figure supplement 1C). Addition of iron, zinc, or manganese did not rescue the growth defects of *ctr1*Δ strains, confirming that the respiratory deficiencies were specifically due to insufficient copper utilization (Figure 6 – figure supplement 1D).

Consistent with the hypomorphic nature of the *H3H113N* mutation, a strain with one WT histone H3 and one containing the *H113N* mutation revealed a partial loss of function in copper utilization that was intermediate to that of strains containing two *H3H113N* or two WT H3 genes (Figure 6F). Furthermore, we tested whether a lower dosage of nucleosomes would also disrupt copper utilization by deleting one of the two copies of the histone H3 and H4 genes (*hht1-hhf1*Δ) in an otherwise WT cell. Accordingly, *hht1-hhf1*Δ displayed decreased cellular H3 and H4 protein content and increased chromatin accessibility by Micrococcal nuclease (MNase) (Figure 6 – figure supplement 2) but did not affect growth in YPD (Figure 6G). Importantly, *hht1-hhf1*Δ increased the requirement of copper in *ctr1*Δ for growth in YPEG medium (Figure 6H).

### Disruptions in copper utilization by H3H113 mutations are not accounted for by changes in chromatin function, cellular copper levels, extracellular metal reduction, or glutathione levels

Our findings are consistent with H3H113 mutants impacting cuprous ion availability, but we considered whether potential disruptions in chromatin structure or gene regulation account for the copper utilization defects as opposed to direct interactions of the histones with copper ions. Deletion of Asf1, a histone chaperone that facilitates nucleosome assembly in part through interaction with H3H113 (Adkins et al., 2004; Agez et al., 2007), or the *H3H113N* mutation resulted in minimal disruption of chromatin accessbility compared to WT as assessed by MNase digestion (Figure 6 – figure supplement 3A). Importantly, any potential disruption of chromatin structure did not account for the increased requirement for copper, as *asf1*Δ did not phenocopy the H3H113 mutants in respiratory media (Figure 6 – figure supplement 3B). Global gene expression patterns were similar between WT and *H3^H113N^* strains in respiratory media (Figure 6 – figure supplement 3C), with comparable upregulation of genes involved in the electron transport chain, tricarboxylic acid cycle, and copper and iron regulation (Figure 6 – figure supplement 3D and E).

We also considered whether changes in the total availability of copper ions could account for the defect in *H3H113N* as opposed to impacts on just the cuprous species. As was the case in fermentative YPD medium (Figure 5C), total intracellular levels of copper were not lower in *H3^H113N^* compared to WT in respiratory YPEG medium, although a slight decrease in iron in YPEG was observed (Figure 6 – figure supplement 4A). Addition of excess copper in YPEG, which rescued mitochondrial respiration in *ctr1*Δ but not in *H3^H113N^ctr1*Δ (Figure 6 – figure supplement 1B), increased intracellular levels of copper similarly in both strains (Figure 6 – figure supplement 4A). These findings indicate that the defect in copper utilization in *H3^H113N^ctr1*Δ was not due to deficient copper uptake through the remaining minor copper importers, such as Fet4 (Hassett et al., 2000). Lastly, inefficient recovery of *ctr1*Δ in the context of *H3H113N* was not due to increased sequestration of copper ions by the metallothionein Cup1 since loss of Cup1 (*cup1^F8stop^*) had no effect on the requirement for copper in *H3^H113N^ctr1*Δ (Figure 6 – figure supplement 4B).

We next asked whether the known extracellular metalloreductases *FRE1* and *FRE2* (Hassett and Kosman, 1995) might increase the copper requirement similar to the H3H113 mutations. Deletion of both FRE genes did not phenocopy the H3H113 mutations, however, (Figure 6 – figure supplement 5A), indicating that an inability to maintain intracellular rather than extracellular cuprous ions results in copper-dependent respiratory deficiency. Furthermore, we examined how the H3H113 mutations would compare to the effect of decreasing cellular levels of glutathione (GSH), which participates in cellular copper metabolism (Freedman et al., 1989). Decreasing GSH abundance, by deleting the gamma glutamylcysteine synthetase gene (*GSH1*), increased the amount of copper required to rescue *ctr1*Δ in the context of *gsh1*Δ but to a lesser extent than the *H3H113N* mutation (Figure 6 – figure supplement 5B). Combining *H3H113N* with *gsh1*Δ caused an even greater defect in copper utilization (Figure 6 – figure supplement 5B), suggesting that the histones impact cuprous ion utilization in a different manner than GSH.

### Histone H3 is required for efficient utilization of copper for Sod1 function

We next considered whether the impact of histone H3 on cuprous ion availability and copper-dependent function extends to the yeast Sod1. An in-gel superoxide dismutase assay (Leitch et al., 2009) demonstrated that total Sod1 activity was ∼40% less in *H3^H113Y^* compared to WT when grown in rich medium, which was recovered by addition of 50 µM copper (Figure 7A and Figure 7 – figure supplement 1A). *H3H113N* also decreased Sod1 activity by ∼20% compared to WT (Figure 7B). Deletion of *CTR1* reduced Sod1 activity and formation of its internal disulfide bond, which are both dependent on the presence of cuprous ions and on the copper chaperone for Sod1, Ccs1 (Furukawa et al., 2004). Combining *ctr1*Δ with *H3H113N* further decreased the internal disulfide bond and substantially lowered Sod1 activity to ∼10% of WT. Both Sod1 activity and its disulfide bond were restored to wildtype levels by addition of excess exogenous copper (Figure 7B).

**Figure 7.**
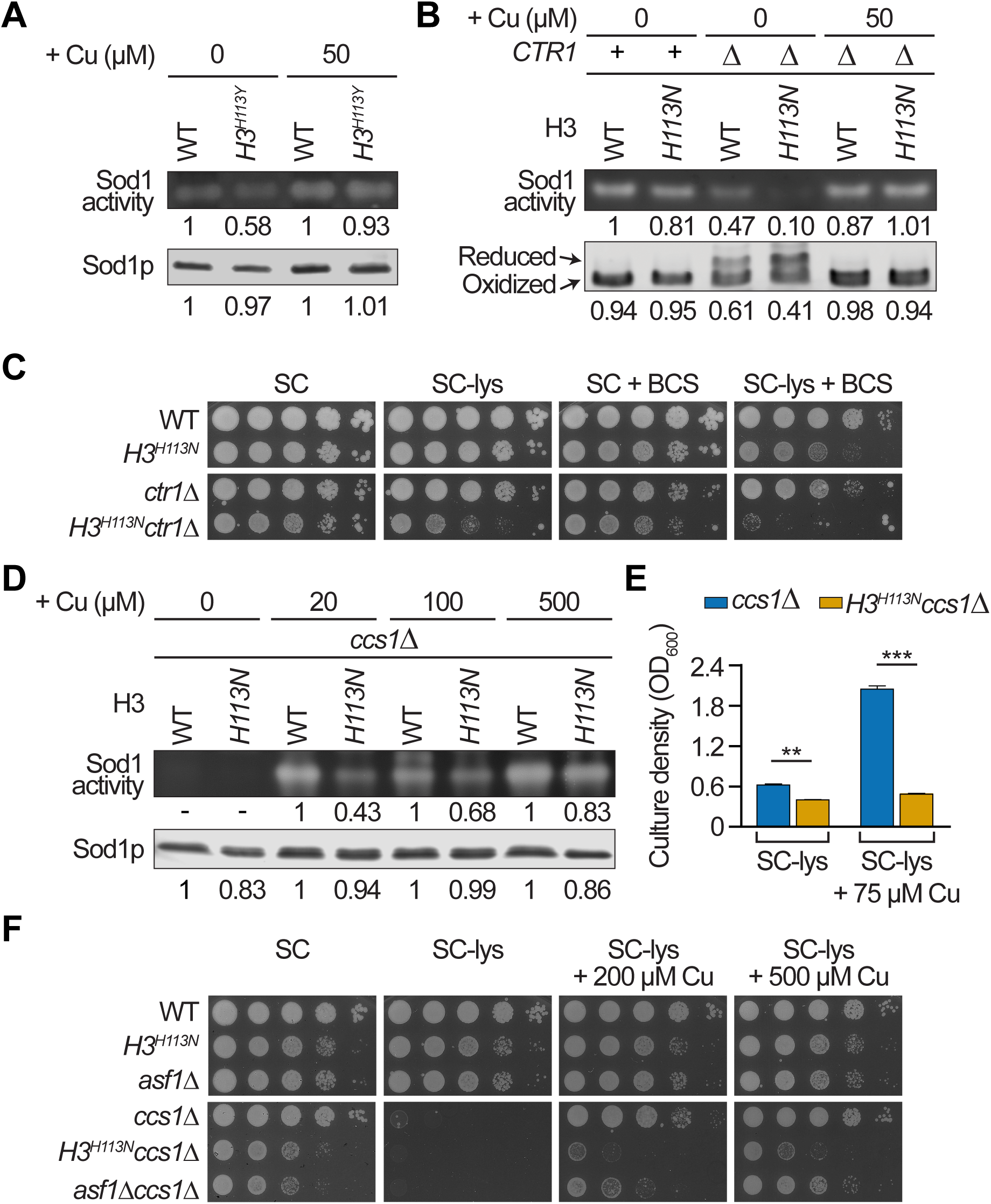
*H3^H113N^* is deficient in utilizing copper for Sod1 activity and function. **(A)** Sod1 in-gel activity assay (top) and corresponding Sod1p western blot (bottom) for cells grown in SC medium with the indicated amounts of additional CuSO_4_. Baseline copper concentration in SC medium is ∼0.16 µM. Numbers are relative signal intensities for each pair of bands. Gel and blot shown are representative of four independent experiments. **(B)** Same as (A), except paired with an Sod1 disulfide bond assay (bottom). Numbers below the blot are relative amount of oxidized Sod1 compared to the total Sod1 protein. Bands shown are representative of three independent experiments. **(C)** Spot test assays on SC or SC-lys plates containing 1 mM and 10 µM BCS, respectively. **(D)** Same as (A) except for cells grown in minimal medium. Baseline copper concentration in minimal medium is ∼0.25 µM. Gel and blot shown are representative of three independent experiments. **(E)** Growth assay in the indicated liquid media shown as mean OD_600_ ± SD after 24 hrs from three independent experiments. **(F)** Same as (C). **P≤0.01, ***P<0.001.

Loss of function of *SOD1* or *CCS1* causes lysine auxotrophy in yeast when grown in the presence of oxygen (Lin and Culotta, 1996). Consistent with this, *H3^H113N^ctr1*Δ exhibited a growth defect in lysine deficient conditions (Figure 7C). Further limiting copper abundance by addition of the copper chelator BCS rendered *H3^H113N^ctr1*Δ, but not *ctr1*Δ, auxotrophic for lysine even in the presence of wildtype *CCS1* and *SOD1* genes (Figure 7C). Together, these data are consistent with a deficiency in copper utilization for Sod1 function when H3H113 is mutated.

We also assessed the ability of cells to restore copper-dependent Sod1 function starting from a state of low activity due to low delivery of copper ions. Copper delivery to Sod1 was first diminished by deleting *CCS1*, which significantly decreased Sod1 activity (Figure 7D). Sod1 activity in the absence of Ccs1 was increasingly restored by addition of excess copper in rich medium (Figure 7D). Importantly, the *H3H113N* mutation increased the amount of copper required to restore Sod1 activity in *ccs1*Δ (Figure 7D). Correspondingly, the lysine auxotrophy of *ccs1*Δ was rescued by addition of excess exogenous copper (Figure 7E). However, substantially more copper was required to rescue *ccs1*Δ in the context of the *H3H113N* mutation (Figure 7E). Interestingly, unlike the effect of *H3H113N*, deletion of *CTR1* did not further increase the requirement of *ccs1*Δ for exogenous copper (Figure 7 – figure supplement 1B), suggesting that histone H3 contributes to copper utilization in a qualitatively different manner than Ctr1. Excess copper did not rescue the lysine auxotrophy of *sod1*Δ strains, confirming that Sod1 is required for copper-dependent lysine prototrophy (Figure 7 – figure supplement 1C). Hypoxia rescued the lysine auxotrophy of *ccs1*Δ and *sod1*Δ in both WT and *H3^H113N^* (Figure 7 – figure supplement 1D), indicating that the copper utilization defect for lysine prototrophy in *H3H113N* strains only manifests when Sod1 function is required.

The *H3H113N* mutation specifically disrupts copper utilization as addition of manganese, zinc, or iron did not rescue *ccs1*Δ lysine auxotrophy (Figure 7 – supplement 1E). The increased copper requirement of *H3^H113N^ccs1*Δ was not due to differences in Cup1-dependent copper sequestration (Figure 7 – figure supplement 2A) or intracellular copper and iron levels (Figure 7 – figure supplement 2B and C). As with respiratory growth, deletion of *ASF1* had no effect on the copper-dependent rescue of *ccs1*Δ (Figure 7F). Global gene expression also did not account for the defective copper utilization as *ccs1*Δ and *H3^H113N^ccs1*Δ displayed similar patterns (Figure 7 – figure supplement 2D), including genes involved in lysine biosynthesis, copper homeostasis, and antioxidant defense (Figure 7 – figure supplement 2E and F). Altogether, our findings reveal that the H3H113 residue is also important for copper utilization by Sod1. The significant impacts of H3H113 mutations in at least two separate copper-dependent processes support the model that the interaction of copper ions and the putative cupric reductase function of the yeast histone H3 impact cuprous ion availability in the cell.

### The *H3A110C* mutation enhances copper utilization for respiratory growth

The ability of H3C110 to enhance copper reductase activity of the yeast tetramer *in vitro* suggested that the *H3A110C* mutation should enhance cuprous ion availability and copper utilization *in vivo*. We generated the yeast *H3^A110C^* strain, with the *H3A110C* mutation in both *HHT1* and *HHT2* genes in their chromosomal loci. This strain grew similarly to WT cells in rich respiratory medium (Figure 8A). However, in contrast to the effects of the H3H113 mutations, the *H3A110C* mutation enhanced the ability of exogenous copper to restore the growth of *ctr1*Δ in YPEG, consistent with more efficient copper utilization for mitochondrial respiration (Figure 8B). We reasoned that increased provision of cuprous ions, due to the enhanced cupric ion reduction caused by the *H3A110C* mutation, may mitigate the depletion of cellular GSH. Indeed, *H3^A110C^gsh1*Δ grew better than *gsh1*Δ in copper-depleted respiratory conditions (Figure 8C) with a greater O_2_ consumption rate (Figure 8D). Furthermore, *H3^A110C^* exhibited a growth advantage upon addition of a toxic but sub-lethal amount of potassium cyanide (KCN), a potent inhibitor of the copper-dependent cytochrome *c* oxidase independently of GSH levels, especially upon addition of exogenous copper (Figure 8E and F). These results highlight the advantage in copper utilization conferred by the *H3A110C* mutation. Moreover, the presence of the *H3A110C* mutation significantly rescued the copper utilization defect of the *H3^H113N^ctr1*Δ strain (Figure 8G). Lastly, the presence of the *H3A110C* mutation rescued the lethality of the *H3H113A* mutation (Figure 8H), corroborating the opposing effects of the H3A110 and H3H113 mutations both *in vivo* and *in vitro*. These findings further support that histone H3 contributes to cellular copper utilization, likely through the maintenance of the cuprous ion pool via catalytic conversion of cupric to cuprous species.

**Figure 8.**
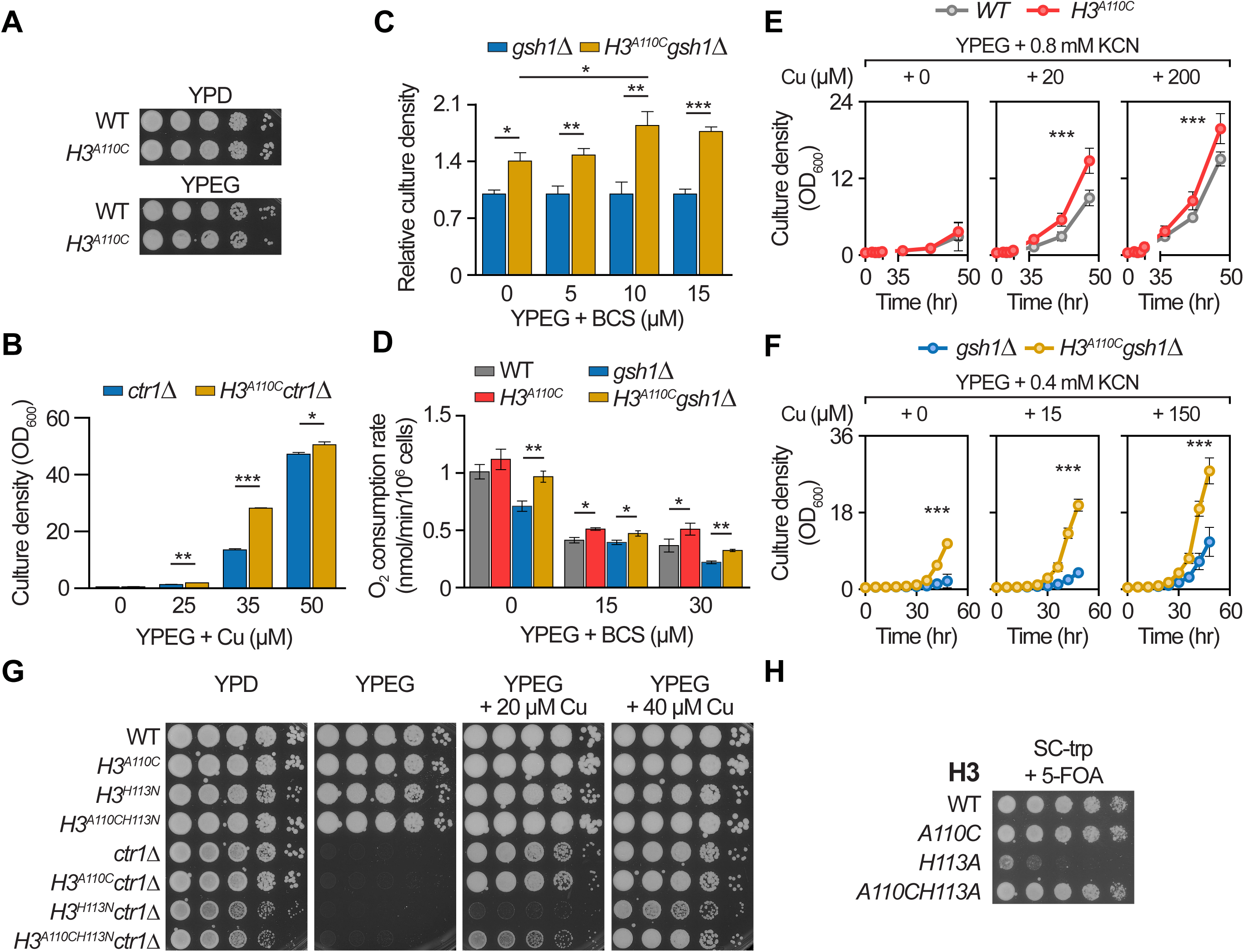
The *H3A110C* mutation enhances copper utilization for respiratory growth in *S. cerevisiae*. **(A)** Spot test assays in fermentative (YPD) or respiratory media (YPEG). **(B)** Growth after 48 hrs in liquid YPEG with increasing amounts of CuSO_4._ Bars show mean OD_600_ ± SD from three independent experiments done concurrently with the same batch of media. **(C)** Growth after 48 hrs in liquid YPEG with increasing amounts of BCS. Bars show relative OD_600_ compared to the *gsh1*Δ strain ± SD from three independent experiments. **(D)** Oxygen consumption assays of cells grown in the indicated media for 12 hrs. Bar graphs show means ± SD of three independent experiments done concurrently with the same batch of media and scaled to relative mitochondrial DNA contents (data not shown). **(E)** Growth curves of the indicated strains in YPEG with potassium cyanide (KCN) and increasing amounts of CuSO_4_ in each panel, as indicated. Line graphs show means at each time point ± SD from three independent experiments done concurrently with the same batch of media. *** indicates significant (P≤0.001) difference at multiple time points. Note that minimal growth occurred before 36 hrs. **(F)** Same as (E). **(G)** Spot test assays in YPD or YPEG with the indicated amounts of CuSO_4_. **(H)** Plasmid shuffle assay with strains harboring WT H3 on a URA3 plasmid and the indicated H3 gene on a TRP1 plasmid. 5-Fluoro-orotic acid (5-FOA) is lethal to cells bearing the URA3 plasmid, and therefore selects for cells that maintain only the indicated H3 plasmid. *P<0.05, **P≤0.01, ***P≤0.001.

## Discussion

Since their identification, histones have been considered to serve as packaging and regulatory proteins for the eukaryotic genome. We now reveal the eukaryotic histone H3-H4 tetramer as a cupric reductase that provides cuprous ions for cellular and mitochondrial biology. The enzymatic activity of the H3-H4 tetramer suggests the protein complex has novel features, including a catalytic site, that were previously unsuspected. The structural features, unusual evolutionary conservation, and our biochemical data suggest that a likely site for the catalytic pocket is at the interface of the two H3 proteins. Subsequent studies should determine the mechanisms by which the residues in this region, including H113 and C110, contribute to direct cupric ion binding, increasing its reduction potential, and promoting electron transfer (Liu et al., 2014). The H3-H3’ interface forms *in vivo* only during nucleosome assembly on DNA and can remain accessible by molecules in the surrounding environment (Feinstein and Moudrianakis, 1986). Therefore, the commencement of enzymatic activity would be inextricably coupled to the protection of DNA as it wraps on the outer surface of the nucleosome away from the putative binding site. Such a coupling may be a beneficial adaptation in species that require provision of reduced copper, as it would also decrease the potentially coinciding damage due to copper.

How the tetramer extracts electrons from a reducing co-factor(s) is unclear, as is the location at which this occurs. Electrons could transfer directly to the tetramer-bound copper from co-factors or through the protein in a mechanism involving a chain of residues in transient redox states (Lucas et al., 2011; Williams et al., 1997). In this regard, whether the sequence differences between the histone H3 variants or post-translational modifications of the histones influence the kinetic properties or co-factor requirements of the H3-H4 tetramer enzyme activity remain interesting and open questions.

Since cellular copper ions are mostly bound by proteins (Rae et al., 1999), copper is likely delivered to and taken away from histones through as-yet-to-be-identified chaperones. Copper is present in the nucleus but the dynamics of its entry and exit or its function(s) are unclear (McRae et al., 2013; Yang et al., 2005). Interestingly, when cellular copper efflux is compromised such as in Wilson disease, copper disproportionately accumulates in the nucleus, reaching millimolar concentrations (Burkhead et al., 2011; Huster et al., 2006), albeit likely bound to chaperones. Considering the importance of copper to processes as diverse as tissue integrity, methylation cycle, iron homeostasis, and melanin and neurotransmitter syntheses in mammals (Finney et al., 2014; Nevitt et al., 2012; Winston and Jaiser, 2008), the enzymatic activity of histones could have wide-ranging effects at molecular, cellular, and tissue levels with consequences for organismal physiology and disease.

The oxidoreductase function of the H3-H4 tetramer, the ancient structural form of histones, could provide a reasonable explanation for why the archaeal ancestor of the eukaryotes possessed histone tetramers. This enzymatic activity, together with an associated intracellular Cu^1+^ transit system, could have facilitated the provision of cuprous ions to the nascent mitochondria to maintain a functioning ETC thereby intimately linking histones to the successful emergence of our common eukaryotic ancestor.

## Materials and Methods

Statistical methods were not used to determine sample size. The investigators were not blinded to the identity of strains and samples in experiments and analysis.

### Strains and general growth conditions

*Saccharomyces cerevisiae* strains used in this study are based on the BY4741 (S288C background, MATa) (Brachmann et al., 1998) and RMY102 (MATa) (Mann and Grunstein, 1992) strains and are listed in Table 1. Strains based on BY4741 were maintained on YPD (1% Yeast extract, 2% Peptone, 2% Glucose) plates, and which were supplemented with 20 µM CuSO_4_ for strains lacking *CTR1*. Furthermore, strains lacking *CTR1* were used for experiments within three passages to avoid emergence of potential mutants with enhanced ability for acquisition of copper through alternative routes. Strains based on RMY102 were maintained on SG-trp-ura (Synthetic complete medium, lacking tryptophan and uracil, and with 2% Galactose) plates. All strains were grown at 30°C in all experiments as described below. Bacteria for histone expression were BL21(DE3)pLysS and BL21(DE3) cells (Agilent). Transformed bacteria were grown on 2xTY media (1.6% Bacto Tryptone, 1% Yeast Extract, 0.5% NaCl) supplemented with 100 μg/mL ampicillin.

**Table 1.** Yeast strains used in this study.

| Strain name | Mutant short name | Genotype | Reference |
| --- | --- | --- | --- |
| BY4741 | parental | *MATa his3Δ1 leu2Δ0 met15Δ0 ura3Δ0* | Brachmann et al., 1998 |
| OCY1131 | WT | *MATa his3Δ1 leu2Δ0 met15Δ0 ura3Δ0, (hht2Δ::ura3)Δ::HHT2* | This study |
| OCY1136 | *H3<sup>H113N</sup>* | *MATa his3Δ1 leu2Δ0 met15Δ0 ura3Δ0, (hht2Δ::ura3)Δ::hht2-H113N hht1-H113N* | This study |
| OCY1133 | *p<sup>(GAL1)</sup>-HHT1* | *MATa his3Δ1 leu2Δ0 met15Δ0 ura3Δ0, (hht2Δ::ura3)Δ::HHT2, KanMX6-PGAL1-HHT1* | This study |
| OCY1134 | *hht2<sup>H113N</sup>p<sup>(GAL1)</sup>-HHT1* | *MATa his3Δ1 leu2Δ0 met15Δ0 ura3Δ0, (hht2Δ::ura3)Δ::hht2-H113N, KanMX6-PGAL1-HHT1* | This study |
| NAY609 | *H3<sup>H113Y</sup>* | *MATa his3Δ1 leu2Δ0 met15Δ0 ura3Δ0, (hht2Δ::ura3)Δ::HHT2, hht2-H113Y, hht1-H113Y* | This study |
| NAY616 | *H3<sup>A110C</sup>* | *MATa his3Δ1 leu2Δ0 met15Δ0 ura3Δ0, (hht2Δ::ura3)Δ::HHT2, hht2-A110C, hht1-A110C* | This study |
| OCY2951 | *hht2<sup>H113Y</sup>p<sup>(GAL1)</sup>-HHT1* | *MATa his3Δ1 leu2Δ0 met15Δ0 ura3Δ0, (hht2Δ::ura3)Δ::HHT2, hht2-H113Y, KanMX6-PGAL1-HHT1* | This study |
| NAY536 | *asf1Δ* | *MATa his3Δ1 leu2Δ0 met15Δ0 ura3Δ0, (hht2::ura3)::HHT2, asf1Δ::HphMX4* | This study |
| OCY1861 | *ctr1Δ* | *MATa his3Δ1 leu2Δ0 met15Δ0 ura3Δ0, (hht2Δ::ura3)Δ::HHT2, ctr1Δ::HphMX4* | This study |
| OCY1862 | *H3<sup>H113N</sup> ctr1Δ* | *MATa his3Δ1 leu2Δ0 met15Δ0 ura3Δ0, (hht2Δ::ura3)Δ::hht2-H113N hht1-H113N, ctr1Δ::HphMX4* | This study |
| OCY2141 | *asf1Δ ctr1Δ* | *MATa his3Δ1 leu2Δ0 met15Δ0 ura3Δ0, (hht2Δ::ura3)Δ::HHT2, asf1Δ::HphMX4, ctr1Δ::His3MX6* | This study |
| OCY2251 | *mac1Δ* | *MATa his3Δ1 leu2Δ0 met15Δ0 ura3Δ0, (hht2Δ::ura3)Δ::HHT2, mac1Δ::His3MX6* | This study |
| OCY2252 | *H3<sup>H113N</sup> mac1Δ* | *MATa his3Δ1 leu2Δ0 met15Δ0 ura3Δ0, (hht2Δ::ura3)Δ::hht2-H113N hht1-H113N, mac1Δ::His3MX6* | This study |
| OCY2621 | *ctr1Δ* | *MATa his3Δ1 leu2Δ0 met15Δ0 ura3Δ0, (hht2Δ::ura3)Δ::HHT2, ctr1Δ* | This study |
| OCY2622 | *H3<sup>H113N</sup> ctr1Δ* | *MATa his3Δ1 leu2Δ0 met15Δ0 ura3Δ0, (hht2Δ::ura3)Δ::hht2-H113N hht1-H113N, ctr1Δ* | This study |
| OCY2623 | *H3<sup>H113Y</sup> ctr1Δ* | *MATa his3Δ1 leu2Δ0 met15Δ0 ura3Δ0, (hht2Δ::ura3)Δ::HHT2, hht2-H113Y, hht1-H113Y, ctr1Δ* | This study |
| OCY1981 | *cup1<sup>F8stop</sup>* | *MATa his3Δ1 leu2Δ0 met15Δ0 ura3Δ0, (hht2Δ::ura3)Δ::HHT2, cup1-F8stop* | This study |
| OCY1983 | *cup1<sup>F8stop</sup>H3<sup>H113N</sup>* | *MATa his3Δ1 leu2Δ0 met15Δ0 ura3Δ0, (hht2Δ::ura3)Δ::HHT2, cup1-F8stop, hht1-H113N hht2-H113N* | This study |
| OCY2181 | *cup1<sup>F8stop</sup>ctr1Δ* | *MATa his3Δ1 leu2Δ0 met15Δ0 ura3Δ0, (hht2Δ::ura3)Δ::HHT2, cup1-F8stop, ctr1Δ::His3MX6* | This study |
| OCY2182 | *cup1<sup>F8stop</sup>H3<sup>H113N</sup>ctr1Δ* | *MATa his3Δ1 leu2Δ0 met15Δ0 ura3Δ0, (hht2Δ::ura3)Δ::HHT2, cup1-F8stop, hht1-H113N hht2-H113N, ctr1Δ::His3MX6* | This study |
| OCY1132 | *hht2<sup>H113N</sup>* | *MATa his3Δ1 leu2Δ0 met15Δ0 ura3Δ0, (hht2Δ::ura3)Δ::hht2-H113N* | This study |
| OCY2628 | *hht2<sup>H113N</sup>ctr1Δ* | *MATa his3Δ1 leu2Δ0 met15Δ0 ura3Δ0, (hht2Δ::ura3)Δ::hht2-H113N, ctr1Δ* | This study |
| NAY553 | *ccs1Δ* | *MATa his3Δ1 leu2Δ0 met15Δ0 ura3Δ0, (hht2::ura3)::HHT2, ccs1Δ::KanMX6* | This study |
| NAY554 | *H3<sup>H113N</sup> ccs1Δ* | *MATa his3Δ1 leu2Δ0 met15Δ0 ura3Δ0, (hht2::ura3)::hht2-H113N hht1-H113N, ccs1Δ::KanMX6* | This study |
| NAY597 | *asf1Δ ccs1Δ* | *MATa his3Δ1 leu2Δ0 met15Δ0 ura3Δ0, (hht2Δ::ura3)Δ::HHT2, asf1Δ::HphMX4, ccs1Δ::His3MX6* | This study |
| NAY551 | *sod1Δ* | *MATa his3Δ1 leu2Δ0 met15Δ0 ura3Δ0, (hht2::ura3)::HHT2, sod1Δ::HphMX4* | This study |
| NAY552 | *H3<sup>H113N</sup> sod1Δ* | *MATa his3Δ1 leu2Δ0 met15Δ0 ura3Δ0, (hht2::ura3)::hht2-H113N hht1-H113N, sod1Δ::HphMX4* | This study |
| OCY2211 | *cup1<sup>F8stop</sup>ccs1Δ* | *MATa his3Δ1 leu2Δ0 met15Δ0 ura3Δ0, (hht2Δ::ura3)Δ::HHT2, cup1-F8stop, ccs1Δ::KanMX6* | This study |
| OCY2212 | *cup1<sup>F8stop</sup>H3<sup>H113N</sup>ccs1Δ* | *MATa his3Δ1 leu2Δ0 met15Δ0 ura3Δ0, (hht2Δ::ura3)Δ::HHT2, cup1-F8stop, hht1-H113N hht2-H113N, ccs1Δ::KanMX6* | This study |
| OCY1135 | *hht1-hhf1Δ* | *MATa his3Δ1 leu2Δ0 met15Δ0 ura3Δ0, (hht2Δ::ura3)Δ::HHT2, hht1-hhf1Δ::HphMX4* | This study |
| OCY2421 | *ctr1Δ* | *MATa his3Δ1 leu2Δ0 met15Δ0 ura3Δ0, (hht2Δ::ura3)Δ::HHT2, ctr1Δ::His3MX6* | This study |
| NAY564 | *hht1-hhf1Δctr1Δ* | *MATa his3Δ1 leu2Δ0 met15Δ0 ura3Δ0, (hht2Δ::ura3)Δ::HHT2, hht1-hhf1Δ::HphMX4, ctr1Δ::His3MX6* | This study |
| OCY2781 | *fre1Δfre2Δ* | *MATa his3Δ1 leu2Δ0 met15Δ0 ura3Δ0 (hht2Δ::ura3)Δ::HHT2, fre1Δ::KanMX6, fre2Δ::His3MX6* | This study |
| OCY2791 | *fre1Δfre2Δctr1Δ* | *MATa his3Δ1 leu2Δ0 met15Δ0 ura3Δ0 (hht2Δ::ura3)Δ::HHT2, fre1Δ::KanMX6, fre2Δ::His3MX6, ctr1Δ* | This study |
| OCY2371 | *gsh1Δ* | *MATa his3Δ1 leu2Δ0 met15Δ0 ura3Δ0, (hht2Δ::ura3)Δ::HHT2, gsh1Δ::KanMX6* | This study |
| OCY2372 | *H3<sup>H113N</sup>gsh1Δ* | *MATa his3Δ1 leu2Δ0 met15Δ0 ura3Δ0, (hht2Δ::ura3)Δ::hht2-H113N hht1-H113N, gsh1Δ::KanMX6* | This study |
| OCY2373 | *H3<sup>A110C</sup>gsh1Δ* | *MATa his3Δ1 leu2Δ0 met15Δ0 ura3Δ0, (hht2Δ::ura3)Δ::hht2-A110C hht1-A110C, gsh1Δ::KanMX6* | This study |
| OCY2626 | *H3<sup>A110C</sup>ctr1Δ* | *MATa his3Δ1 leu2Δ0 met15Δ0 ura3Δ0, (hht2Δ::ura3)Δ::HHT2, hht2-A110C, hht1-A110C, ctr1Δ* | This study |
| OCY2671 | *H3<sup>A110CH113N</sup>* | *MATa his3Δ1 leu2Δ0 met15Δ0 ura3Δ0 (hht2Δ::ura3)Δ::HHT2, hht2-A110CH113N, hht1-A110CH113N* | This study |
| OCY2627 | *H3<sup>A110CH113N</sup>ctr1Δ* | *MATa his3Δ1 leu2Δ0 met15Δ0 ura3Δ0 (hht2Δ::ura3)Δ::HHT2, hht2-A110CH113N, hht1-A110CH113N, ctr1Δ* | This study |
| RMY102 | parental | *MATa ade2-101 his3-Δ200 lys2-801 trp1Δ901 ura3-52 (hht1-hhf1)Δ::LEU2 (hht2-hhf2)Δ::HIS3 pRM102[CEN4 ARS1 URA3 pGAL1-HHT2-pGAL10-HHF2]* | Mann & Grunstein, 1998 |
| NAY702 | WT | *MATa ade2-101 his3-Δ200 lys2-801 trp1Δ901 ura3-52 (hht1-hhf1)Δ::LEU2 (hht2-hhf2)Δ::HIS3 pRM102[CEN4 ARS1 URA3 pGAL1-HHT2-pGAL10-HHF2] pRM200[CEN4 ARS1 TRP1 HHT2 HHF2]* | This study |
| NAY700 | *H3-H113A* | *MATa ade2-101 his3-Δ200 lys2-801 trp1Δ901 ura3-52 (hht1-hhf1)Δ::LEU2 (hht2-hhf2)Δ::HIS3 pRM102[CEN4 ARS1 URA3 pGAL1-HHT2-pGAL10-HHF2] pSZ008[CEN4 ARS1 TRP1 hht2-H113A HHF2]* | This study |
| NAY703 | *H3-H113N* | *MATa ade2-101 his3-Δ200 lys2-801 trp1Δ901 ura3-52 (hht1-hhf1)Δ::LEU2 (hht2-hhf2)Δ::HIS3 pRM102[CEN4 ARS1 URA3 pGAL1-HHT2-pGAL10-HHF2] pYX55a[CEN4 ARS1 TRP1 hht2-H113N HHF2]* | This study |
| **NAY701** | *H3-A110C* | *MATa ade2-101 his3-Δ200 lys2-801 trp1Δ901 ura3-52 (hht1-hhf1)Δ::LEU2 (hht2-hhf2)Δ::HIS3 pRM102[CEN4 ARS1 URA3 pGAL1-HHT2-pGAL10-HHF2] pSZ011[CEN4 ARS1 TRP1 hht2-A110C HHF2]* | This study |
| **NAY704** | *H3-A110CH113A* | *MATa ade2-101 his3-Δ200 lys2-801 trp1Δ901 ura3-52 (hht1-hhf1)Δ::LEU2 (hht2-hhf2)Δ::HIS3 pRM102[CEN4 ARS1 URA3 pGAL1-HHT2-pGAL10-HHF2] pNA20a[CEN4 ARS1 TRP1 hht2-A110CH113A HHF2]* | This study |

### Strain generation for yeast experiments

*BY4741 background:* The histone H3 *H113N* mutation was generated in both chromosomal loci (*HHT1* and *HHT2*) in a stepwise and marker-less manner. First, using the *delitto perfetto* approach (Storici and Resnick, 2006), the *H113N* mutation was introduced into the *HHT2* locus. Second, the *H113N* mutation was introduced in *HHT1* using the CRISPR-Cas9 system optimized for *S. cerevisiae* (Ryan et al., 2014). The pCAS plasmid, containing both the Cas9 gene from *Streptococcus pyogenes* and a single guide RNA, was a gift from Jamie Cate (Ryan et al., 2014). The histone H3 *H113Y* and *A110C* mutations were introduced in a similar manner as described but CRISPR-Cas9 was used for both HHT1 and HHT2. Subsequent gene deletions and promoter insertions were generated by standard yeast gene replacement and targeted insertion methodology using selectable marker integration (Goldstein and McCusker, 1999; Longtine et al., 1998). Additionally, *CTR1* was deleted in several strains using the CRISPR-Cas9 targeting system. Successful integrations and deletions were confirmed by PCR. The CUP1 gene was disrupted by introduction of a stop codon at Phe8 at all CUP1 copies using the CRISPR-Cas9 system as described above for generating histone mutants.

*RMY102 background:* The histone H3 *H113N*, *H113A*, and *A110C* mutations were introduced into the *HHT2* gene of the pRM200 plasmid (CEN4/ARS1/TRP1/HHT2-HHF2) (Mann and Grunstein, 1992) by site directed mutagenesis (QuikChange, Agilent). The WT or mutagenized pRM200 plasmids were then introduced into the RMY102 strain by standard yeast transformation procedures, and cells were maintained on media lacking both tryptophan and uracil to ensure the presence of both pRM102 (CEN4/ARS1/URA3/p^GAL1^-HHT2-p^GAL10^-HHF2) and pRM200 plasmids.

### Plasmids for histone expression

Plasmids for expression of *Xenopus laevis* histones are from (Luger et al., 1997). All sequence changes to the plasmid were accomplished by site directed mutagenesis (QuikChange, Agilent). The histone H3 plasmid from this reference contains an H3-G102A mutation, which was corrected to the WT G102. H3C110 was mutated to alanine and H3H113 was mutated to alanine, asparagine or tyrosine. pBL105 and pBL108 plasmids for *Saccaromyces cerevisiae* H3 and H4, respectively, were kindly provided by Bing Li, PhD, UT Southwestern Medical Center. H3A110 was mutated to cysteine and H3H113 was mutated to alanine, asparagine or tyrosine.

### Preparation of solutions and glassware

All glassware was treated with 3.7% hydrochloric acid for 12 hours followed in some cases by 12 hours of 10% nitric acid to remove trace metal contamination. All solutions, buffers, and washes were prepared using Nanopure Diamond (Thermo Fisher) ultra-pure water. Solutions were prepared using Bioultra grade reagents and remaining trace metals removed with Chelex 100 (Sigma). Standard RC dialysis tubing for dialysis (Spectrumlabs.com) was washed twice in 10 mM EDTA (99.99% trace metal basis – Sigma) at 80°C and cooled to room temperature for 16 hours. For yeast media, addition of all components was done without the use of metal spatulas. Media was filtered through 0.2 µm membranes. Fermentative media was either YPD (1% Yeast Extract, 2% Peptone, 2% glucose), SC (synthetic complete medium with 2% glucose, all amino acids, and uracil and adenine), SC lacking lysine, or minimal medium (with 2% glucose and only the auxotrophic metabolites His, Leu, Met, and uracil). Non-fermentative media was YPEG (1% Yeast Extract, 2% Peptone, 3% ethanol, 3% glycerol). Agar media was prepared similarly using acid-washed glassware. Agar media for hypoxic growth were supplemented with 20 µg/ml ergosterol, 0.5% Tween-80 and 0.5% ethanol.

### Histone purification

Histone purification was performed as previously described (Luger et al., 1999), except that purification by size exclusion was omitted and the solubilized inclusion bodies were pre-cleaned by passage over a Hi-Trap Q-Sepharose column (GE healthcare) before loading onto a Hi-Trap SP-Sepharose column (GE healthcare). Histones were eluted by a salt gradient from 200 mM to 600 mM NaCl, as described previously (Luger et al., 1999), and fractions containing pure histones were pooled, dialyzed three times against 4 L of 2 mM β-mercaptoethanol (Sigma) in water, flash frozen and lyophilized.

### Tetramer assembly

H3-H4 tetramer assembly was performed as previously described (Luger et al., 1999) with some adjustments. Briefly, equimolar amounts of H3 and H4 were dissolved in 7 M Guanidinium HCl, 20 mM Tris-HCl pH 7.5 and 10 mM DTT at a concentration of 1-2 mg/mL total protein. Refolding was achieved by 1-2 hr dialysis at 4°C against 1 L of 2 M NaCl, 10 mM Tris-HCl pH 7.5, 1 mM EDTA, 5 mM β-mercaptoethanol (T.E.B.), then 1-2 hr against 1 L of 1 M NaCl, T.E.B., followed by 16 hrs against 0.5 M NaCl, T.E.B., except for *S. cerevisiae* H3-H4 tetramers used for Figure 3A, for which all dialysis steps were performed in 2 M NaCl T.E.B.. The refolded tetramer was centrifuged for 5 min at 13000 rcf at 4°C to eliminate any insoluble particles and purified by size exclusion chromatography on a HiLoad 16/600 Superdex 200 column (GE healthcare) at 1 mL/min in 500mM NaCl, 1 mM Tris-HCl pH7.5. Fractions containing the tetramer were concentrated using Amicon Ultra–15/Ultracel-10K centrifugal filters (Millipore). The flow-through buffer was used as negative control (“buffer”) in the copper reductase assay.

### *In vitro* copper reductase assay

H3-H4 tetramer, or the same volume of control buffer, was resuspended to a final concentration of 1 μM in 100 mM NaCl, 1 mM Neocuproine (Sigma) and 30 μM BioUltra TCEP (Sigma) pH 7.5. The reaction was started by adding a mix of CuCl_2_ and Tricine-HCl, pH 7.5, at final concentrations of 0.0125-0.5 mM and 0.125-5 mM, respectively. Special care was taken to keep the ratio of CuCl_2_: Neocuproine at 1:2 for all reactions. CuCl_2_ reduction using NADPH (Sigma, Roche) as the electron donor was performed using a final concentration of H3-H4 tetramer of 5.8 µM in 150 mM NaOAc pH 7.5, 33 mM NaCl, 1 mM Neocuproine, 10 µM NADPH and started by the addition of 0.5 mM CuCl_2_ / 7.5 mM Tricine-HCl, pH 7.5. To reduce a solution of 0.5 mM Cu^2+^-N-(2-Acetamido)iminodiacetic acid (ADA) (Sigma), 5 uM of H3-H4 tetramer were resuspended in 100 mM NaCl, 0.5 mM bicinchoninic acid (BCA) (Sigma) and 100 µM TCEP. The reaction was started by addition of the substrate. Reduction of 0.5 mM Cu^2+^-Nitrilotriacetic acid (NTA) (Sigma) was performed similarly, except for the addition of 200 µM NADH (Roche) as electron donor instead of TCEP. Absorbance at 448 nm (Neocuproine) or 562 nm (BCA) was measured every 0.5 seconds using a Hewlett-Packard HP8453 diode-array UV/Visible spectrophotometer. Standard curves for each condition were produced using 2 mM β-Mercaptoethanol to ensure complete reduction of Cu^2+^. Linearity of detection was verified and the resulting parameters used to convert the absorbance units into Cu^1+^ molarity.

### Proteinase K treatment of tetramer

Proteinase K (Sigma Aldrich) was dissolved in water at 20mg/mL and buffer exchanged into water using Zeba Spin desalting columns (Thermo Fisher Scientific) to clean the enzyme from impurities. 200 µL of 1 µM tetramer or histone H3 reaction mix was incubated with 100 ng of clean Proteinase K for 15 minutes at room temperature before starting the *in vitro* copper reductase assay described above.

### N-Ethylmaleimide (NEM) treatment of tetramer

10 µM *Xenopus laevis* tetramer was incubated with 1 mM NEM in 10 mM Tris-HCl pH 7.5, 500 mM NaCl for 30 min at room temperature. NEM-treated and untreated tetramers were buffer exchanged into 1 mM Tris-HCl pH 7.5, 500 mM NaCl using Zeba Spin desalting columns, concentrated in Amicon Ultra Centrifugal Filters (Millipore) and reacted in the *in vitro* copper reductase assay described above.

### UV/visible Spectrophotometry

Control buffer of each tetramer was used to zero background absorption. The UV/visible spectra of 50 µM *X. laevis* H3-H4 tetramer in 500 mM NaCl and 5 mM Tris-HCl pH 7.5 was acquired first without addition of Cu^2+^. This was followed by addition of 0.25 or 0.5 molar equivalents of CuSO_4_-Tricine pH 7.5 at a ratio of 1:10. The UV/visible spectra of 500 µM *S. cerevisiae* H3-H4 tetramer in 2 M NaCl and 5 mM Tris-HCl pH 7.5 were analyzed in the same manner. The same analysis was performed with the control buffer in the absence of the tetramer and used as “buffer control”. The Hewlett-Packard HP8453 diode-array UV/Visible spectrophotometer was used for all measurements.

### Immobilized Metal Affinity Chromatography (IMAC)

IMAC HiTrap HP (GE Healthcare) columns were washed with 10 column volumes (CV) of water before loading with 5 CV of 5 mM CuCl_2_. Unbound CuCl_2_ was washed off with 10 CV water and the column was equilibrated with 10 CV of Binding Buffer (20 mM Imidazole, 1 mM Tris-HCl pH 7.5, 500 mM NaCl). 325-360 µg of Tetramer in 1 mM Tris-HCl pH 7.5, 500 mM NaCl were loaded twice and washed with 10 CV Binding Buffer followed by 10 CV Binding Buffer containing 100 mM imidazole. Elution was performed using a 100-300 mM imidazole gradient for *S. cerevisiae* tetramers and a 20-1000 mM imidazole gradient for *X. laevis* tetramers over 10 CV. Tetramer background binding to the unloaded IMAC HiTrap HP was competed off with Binding Buffer containing 100 mM imidazole.

### Liquid culture growth curves

Population doubling times were calculated from cells growing exponentially in YPD or SC media. Total population doublings were calculated by measuring cell densities (OD_600_) in indicated media after up to 48 hours of incubation. For experiments involving potassium cyanide treatment, cells from dense starter cultures were washed twice with 5 mM EDTA pH 8 and once with water just before growth.

### Spot tests

Cells from exponentially-growing cultures were 10-fold serially diluted and 5 µL of cells were spotted on media plates as indicated in the figures. For experiments with the *H113N* mutation in a single copy of the histone H3 gene, cells were washed twice with 5 mM EDTA pH 8 prior to spotting. Cells were incubated at 30°C for up to 7 days and imaged daily. Because of differing growth rates in the various media conditions, images shown in the figures were captured when sufficient growth had occurred and growth differences could be assessed, and this ranged between 2-7 days. Experiments in hypoxic conditions were done using a 2.5 L sealable jar and anaerobic gas generating sachets (AnaeroGen) designed to generate a hypoxic environment (oxygen of <1%). 5-fold serially diluted cells were spotted on agar plates. Media for normoxic and hypoxic conditions was supplemented with ergosterol and Tween-80 to support growth in low-oxygen condition and sealed in the hypoxic jar for 7 days at 30°C before capturing images. For plasmid shuffle spot tests, cells growing in SG medium lacking only tryptophan were 5-fold serially diluted, and spotted on media containing 1 mg/mL 5-Fluoroorotic acid. Cells that survive have lost pRM102, but maintain pRM200.

### Micrococcal nuclease (MNase) digestion

MNase digestion was performed similarly to methods described previously (Rando, 2010). Cells were grown in SC medium at 30°C for ∼2 doublings, fixed with 2% formaldehyde for 30 min at 30°C followed by glycine addition to quench formaldehyde reactivity. 2×10^9^ cells were transferred to new tubes and resuspended in Zymolyase buffer (1 M Sorbitol, 50 mM Tris pH 7.5, 10 mM β-mercaptoethanol) on ice. To initiate cell wall digestion, 2 mg of Zymolyase 100-T was added and cells were incubated at 30°C for 35 min. Spheroplasts were resuspended in MNase digestion buffer (1 M sorbitol, 50 mM NaCl, 10 mM Tris pH 7.5, 5 mM MgCl_2_, 1 mM CaCl_2_, 500 µM spermidine, 0.075% Igepal CA-630, 10 mM BME) and varying amounts of MNase (Sigma) were added. Samples were incubated at 37°C for 20 min before inactivation of MNase by 5X stop solution (5% SDS, 50 mM EDTA). Proteins were then digested by proteinase K (Sigma), and formaldehyde crosslinks reversed, by incubation at 65°C for ∼12 hrs. DNA was purified by standard phenol-chloroform extraction methods and isopropanol precipitation. MNase-digested DNA samples were treated with RNase A (Roche) for 1 hr at 37°C. DNA samples were then purified using the Wizard SV PCR purification kit (Promega) according to the manufacturer’s protocol.

### MNase accessibility analysis

MNase-digested DNA samples were normalized by concentration, and visualized using the DNA ScreenTape assay on the TapeStation 2200 instrument (Agilent), according to the manufacturer’s protocol. Gel images exported with TapeStation Analysis software (Agilent) were analyized using FIJI (ImageJ 1.48k). ScreenTape images shown and quantified are representative examples of three replicate MNase digestions from separately grown cultures of the same cell clones.

### Sample preparation for inductively-coupled plasma mass spectrometry (ICP-MS)

Cells from overnight cultures in SC or YPD media were diluted to OD_600_ = 0.2-0.4 and grown at 30°C for ∼3 doublings, or for 18 and 24 hrs in YPEG and SC-lys, respectively. Cells (4-12 × 10^8^) were collected, and washed twice in 1 mM EDTA to remove cell surface-associated metals and once in water. Cell pellets were stored at −20°C for 1-3 weeks prior to preparation for ICP-MS. Plastic bottles and cylinders used for preparation of solutions for sample digestion were treated with 7-10% ACS grade nitric acid for 5 days at 50°C and rinsed thoroughly with water. Previously-stored cell pellets were overlaid slowly with 70% Optima Grade nitric acid and digested at 65°C for 12-16 hrs. Prior to mass spectrometry, the digested samples were diluted with water to a final concentration of 2% nitric acid and a final volume of 3-5 mL.

### Inductively-coupled plasma mass spectrometry

Total Fe and Cu content was measured by inductively coupled plasma mass spectrometry on the Agilent 8800 ICP-QQQ in MS/MS mode. The most abundant isotopes of iron and copper (i.e. Fe^56^ and Cu^63^) were used to determine the total cellular iron and copper levels. The metal content was determined using an environmental calibration standard (Agilent 5183-4688), which contains 100x higher concentration of Fe compared to Cu, which more accurately reflects biological samples. Every run was calibrated individually, and ^45^Sc or ^89^Y were used as internal standards to compare the calibration with the analyzed samples. All Cu and Fe measurements in our study were within the calibrated linear ranges and above the lower limits of detection, as determined from multiple blank samples. ICP MassHunter software was used for ICP-MS data analysis.

### RNA extraction

For poly-A RNA sequencing analyses, cells were growing exponentially in SC or YPD, or grown for 24 hrs in SC-lys at the time of harvesting. For RT-qPCR analyses, cells were growing exponentially in YPD or were exposed to CuSO_4_ for one hour in YPD at the time of harvesting. In both cases, cells were collected by centrifugation and frozen at −20°C until further processing. RNA was extracted using previously published methods (Schmitt et al., 1990). RNA extracted for subsequent RNA-seq analysis are from two replicate experiments. The exceptions are cells grown in SC which are from three replicates, and cells grown in SC-lys + 75 µM CuSO_4_ which is from one experiment. RNA extracted for subsequent RT-PCR analysis are from four replicate experiments.

### Sample preparation for poly-A RNA sequencing

Prior to preparing RNA-seq libraries for Illumina HiSeq sequencing, contaminating DNA was digested using Turbo DNase (Thermo Fisher). RNA quality was then assessed using the RNA ScreenTape assay on the TapeStation 2200 instrument (Agilent), according to the manufacturer’s protocol. Using TapeStation Analysis software (Agilent), RNA Integrity Number equivalent (RINe) scores were calculated. Only samples with RINe scores greater than 9 (out of 10) were used for sequencing library preparation. RNA-sequencing libraries were then prepared either manually with the KAPA Stranded mRNA-seq library prep kit (KAPA Biosystems), or with automation, using the Illumina TruSeq Stranded mRNA Library Kit for NeoPrep (Illumina). For both approaches, libraries were prepared according to the manufacturer’s protocols. RNA-seq libraries were assessed for correct fragment size and the presence of adapter dimers, using the DNA ScreenTape assay on the TapeStation 2200 instrument (Agilent). Average library sizes of ∼270 bp were observed and deemed correct. Libraries were pooled for multiplexed sequencing and further purified using Agencourt RNAClean XP beads. Total DNA concentration was adjusted to 10 nM for Illumina sequencing.

### mRNA-sequencing and data processing

High throughput sequencing was performed on Illumina’s HiSeq 4000 system, with single-end 50 bp insert reads, and dedicated index reads. Total read count per library ranged from ∼1.5-9 million. De-multiplexed reads, in FASTQ file format, were aligned to the R64-1-1 S288C reference genome assembly (sacCer3), which was downloaded from the UCSC database, using Tophat 2.0.9 (Kim et al., 2013). The “-g 1” parameter was used to limit the number of alignments for each read to one, the top-scoring alignment.

### Gene expression analysis

Gene expression values, in reads per kilobase per million mapped reads (RPKMs), for 6692 annotated open reading frames were calculated using SAMMate 2.7.4 (Xu et al., 2011). Many of the 6692 annotated ORFs are labeled as “dubious” or “putative” in the Saccharomyces Genome Database (SGD). They break down as follows: Dubious open reading frame unlikely to encode a functional protein – 717; Putative protein of unknown function – 316; Retrotransposon TYA Gag and TYB Pol genes – 90; Protein of unknown function – 141; Pseudogenic fragment – 4; Other −20. These 1288 annotated ORFs typically have low expression values and were filtered out of the final analyses. The remaining 5404 ORFs were used for comparisons between groups.

### RNA-sequencing gene sets

Gene sets (e.g. Figure 6 – figure supplement 3D) were constructed by downloading and merging gene ontology term gene lists from AmiGO 2 in March 2017. The “copper homeostasis” and “iron homeostasis” gene sets were further modified by adding or removing genes based on literature review.

### RT-qPCR

Extracted nucleic acids were treated with Turbo DNase (Thermo Fisher) and further purified with TRIzol (Thermo Fisher). Poly-A RNA was isolated using Dynabead oligo dT_25_ magnetic beads (Thermo Fisher) and quantified using the Qubit RNA HS assay kit (Thermo Fisher). Equal amounts of mRNA from the different samples were then used to synthesize cDNA, with random hexamers as primers, using the SuperScript IV first-strand synthesis system (Thermo Fisher). Five percent of each cDNA reaction was used in qPCR reactions prepared with the PowerUP SYBR Green Master Mix (Thermo Fisher) and run on a Stratagene Mx3000P qPCR instrument (Agilent). Primer pairs were designed to amplify regions of the Mac1 targets FRE7, CTR1, IRC7, FRE1, and REE1 (Gross et al., 2000), and the loading control ACT1. Fold changes were based on C_t_ values, normalized to ACT1, and calculated relative to the mean of WT in YPD.

### Oxygen consumption assay

Oxygen consumption rates were measured in whole cells using the Fiber Optic Oxygen Monitor, Model 110 (Instech laboratories Inc., Plymouth Meeting, PA). For experiments in Figure 6A and B, cells growing exponentially in YPD or SC were diluted in either YPEG or SCEG + 60 µM BCS, respectively, with or without additional CuSO_4_, and incubated for 18-24 hrs. Cells were diluted sufficiently in YPEG such that final cell density in respiratory media was less than OD_600_ = 3. For experiments in Figure 6C, cells were diluted to OD_600_ = 1 in YPEG with or without additional CuSO_4_ and incubated for 4 hrs. Note that no growth had occurred in that time period due to the respiratory deficiency of *ctr1*Δ strains. For experiments in Figure 8, cells were diluted in YPEG with or without BCS and grown at 30°C for 12 hrs. For experiments involving Antimycin A treatment in YPEG, cells were incubated for 6 hrs in the presence of antimycin. Cells were collected by centrifugation and pellets concentrated in fresh media for the oxygen consumption measurements. Consumption was recorded for 5 minutes and the rate was determined for a one-minute linear consumption period.

### Mitochondrial DNA quantification

Cells grown as described above for oxygen consumption experiments were collected by centrifugation and lysed by bead beating in Lysis buffer 1 (100 mM NaCl, 1 mM EDTA, 10 mM Tris pH 8). Lysates were then treated with RNase A (Sigma) at 37°C 1 hr in Lysis buffer 2 (Lysis buffer 1 with 0.5% Triton X-100) and subsequently with proteinase K (Sigma) at 56°C for 2 hrs in Lysis buffer 3 (Lysis buffer 1 with 2% Triton X-100 and 1% SDS). DNA was purified by standard phenol-chloroform extraction methods and ethanol precipitation. 0.1 ng of total DNA was used in qPCR reactions prepared with the PowerUP SYBR Green Master Mix (Thermo Fisher) and run on a Stratagene Mx3000P qPCR instrument (Agilent). Primer pairs were designed to amplify regions of the COX3 gene on the mitochondrial genome, and ACT1 on the nuclear genome. Mitochondrial DNA (mtDNA) copy numbers were estimated based on C_t_ values compared to ACT1. Oxygen consumption rates were scaled by the relative difference in mtDNA content.

### Cytochrome *c* Oxidase assay

Cells growing exponentially in YPD were diluted to OD_600_ = 1 in YPEG with or without additional CuSO_4_ and incubated for 4 hrs. Note that no growth had occurred in that time period due to the respiratory deficiency of *ctr1*Δ strains. A mitochondria-enriched subcellular fraction was then prepared from ∼2×10^9^ cells using methods described previously (Gregg et al., 2009), with some modifications. Prior to spheroplasting, cells were washed once with 1 mM EDTA, once with water. Cell wall digestion was accomplished using Zymolyase 100-T (Nacalai Tesque) for 35 min at 30°C. The crude mitochondrial fraction was resuspended in 60 µL of Enzyme buffer (10 mM Tris/HCl pH 7, 250 mM sucrose, 0.5% Tween 20). Cytochrome c oxidase activity was assessed similarly to a method described previously (Spinazzi et al., 2012). 1 mM Cytochrome *c* (from equine heart, Sigma) was reduced by addition of 1 mM sodium hydrosulfite. Full reduction was confirmed by the ratio of absorbance at 550 nm to 565 nm being greater than 10. Reactions were carried out in 100 µL volumes at 30°C with 85 µL of Assay buffer (10 mM Tris/HCl ph 7, 120 mM KCl), 5 µL of reduced cytochrome c (50 µM) and 10 µL of mitochondria-enriched samples at a final concentration of 20 – 50 µg/mL. Cytochrome c oxidation was assessed by measuring absorbance at 550 nm over the course of 5 min using a Hewlett-Packard HP8453 diode-array UV/Visible spectrophotometer, and the rate was determined for a one-minute linear oxidation period.

### Sample preparation for Sod1 assays

Sod1p superoxide dismutase activity was assayed by native-PAGE and in-gel staining with nitrotetrazolium blue (NBT), and the Sod1p disulfide bond was assayed by IAA labeling of thiols and gel mobility shift, based on previously-reported methodology (Leitch et al., 2009). For WT and *ctr1*Δ strains (Figure 7A and B), ∼4×10^8^ exponentially growing cells in SC with or without supplemental CuSO_4_ were collected and resuspended in Lysis Buffer (600 mM sorbitol, 10 mM HEPES pH 7.5, 5 mM EDTA and protease inhibitors) containing 1 mM iodoacetamide (IAA) (Figure 7B only). The lysate was split in two for Sod1 activity assays and assessment of the Sod1 disulfide bond. For *ccs1*Δ strains (Figure 7D), following an overnight growth in SC, cells were diluted to OD_600_ = 1 in minimal media with the indicated amounts of CuSO_4_. Cells were incubated at 30°C for 2 hrs, at which point ∼2×10^9^ cells were collected and resuspended in Lysis Buffer. Cells were lysed with glass beads. Lysates were prepared for both native-PAGE and SDS-PAGE with either native sample buffer (12.5% glycerol, 31.25 mM Tris pH 6.8, 0.005% bromophenol blue, 5 mM EDTA final concentrations) or SDS sample buffer (5% glycerol, 40 mM Tris pH 6.8, 1% SDS, 0.005% bromophenol blue) with (Sod1p content) or without (disulfide bond) 50 mM DTT, respectively.

### In-gel Sod1 activity assay

For native-PAGE, ∼10 μg (WT, *H3^H113N^*, and *H3^H113Y^*), ∼20 μg (*ctr1*Δ and *H3^H113N^ctr1*Δ) or ∼100 μg (*ccs1*Δ and *H3^H113N^ccs1*Δ) of total protein was run on a 10% acrylamide gel for 2.5 hrs with 1 mM EDTA in the running buffer. Gels were then sequentially incubated in phosphate buffer (50 mM K_2_HPO_4_ pH 7.8, 1 mM EDTA), 0.48 mM NBT (Sigma) in phosphate buffer, and 30 μM riboflavin (Sigma) and 0.02% Tetramethylethylenediamine (TEMED; ThermoFisher) in phosphate buffer at room temperature. Finally, gels were exposed to bright light and scanned using an Epson document scanner. Signal intensities of the major SOD1 band were then quantified using FIJI (ImageJ 1.48k). For the analysis in Figure 7 – figure supplement 1A, signal intensities from four separate gels were normalized based on the average signal intensity in each gel. Relative signal intensities, compared to the WT sample, were then calculated based on the normalized intensities.

### Sod1p Western Blotting and Sod1 internal disulfide bond measurement

For western blots to determine Sod1p content, 10 µg of total protein content was run on 10% SDS acrylamide gels. Western blots were performed using a primary antibody against Sod1p, a gift from Valeria Culotta (JH764) (Leitch et al., 2009). The antibody was diluted 1:5000 in Odyssey blocking buffer (LI-COR biotechnology) and blots were incubated at room temperature for 1.5 hrs. Signal quantifications were obtained using the Odyssey infrared fluorescence imaging system (LI-COR biotechnology) and Image Studio Lite software. For western blots to determine the amount of disulfide bond, 10 μg (WT, *H3^H113N^*, and *H3^H113Y^*) or 15 μg (*ctr1*Δ and *H3^H113N^ctr1*Δ) of total protein content was run on 18% SDS acrylamide gels. After electrophoresis, gels were incubated for 45 min with SDS running buffer containing 5% β-mercaptoethanol at room temperature. Western blotting for Sod1p, and signal quantification were performed as described above, except the primary antibody was diluted 1:2500.

### SDS-PAGE and Western Blotting for histone content

For assessment of histone protein content, whole cell protein extracts were prepared from 5×10^8^ cells growing exponentially in YPD. Cell pellets were resuspended in 0.2 M NaOH and incubated at room temperature for 15 minutes. Cells were then pelleted and resuspended in SDS-PAGE loading buffer (50 mM Tris-HCl pH 6.8, 2% SDS, 10% glycerol, 1% β-mercaptoethanol) and boiled for 5 minutes. For western blots, 4 µg of total protein was run on 15% SDS acrylamide gels. Primary antibodies to histone H3 (Active Motif, 39163) and H4 (Abcam, 10158) were diluted in Odyssey blocking buffer (LI-COR biotechnology) at 1:5000 and 1:800, respectively, and blots were incubated in diluted antibody at 4°C overnight. Signal quantification was performed as described above.

### Statistical analyses

The number of experimental replicates (n), and the observed significance levels are indicated in figure legends. All statistical analyses were performed using Graphpad Prism 5 or 7, unless otherwise stated. Significance values for pair-wise comparisons of doubling times were calculated using the Mann-Whitney test. Significance values for comparisons of the oxygen consumption rates in Figure 6 – figure supplement 1A were calculated using the Holm-Sidak method for multiple t-tests, with alpha = 0.05. Significance values for pair-wise comparisons of population doublings in Figure 6 – figure supplement 1B, oxygen consumption rates in Figures 6A-C and 8D, cytochrome *c* oxidase activity in Figure 6D, culture densities in Figures 6H, 7E, 8B, and 8C, relative Sod1 activity assay signal intensities in Figure 7 – figure supplement 1A, and gene expression fold changes in Figure 5C were calculated using unpaired Welch’s t-test. Significance values for comparisons of culture densities in Figures 8E and F were calculated using a mixed model Two-Way ANOVA with Bonferroni post-hoc test for pair-wise comparisons. Statistical testing for iron and copper measurements was done using unpaired Student’s t-tests. For gene expression data, we averaged RPKM values from replicates and used mean values for calculation of global gene expression correlations (e.g. Figure 5 – figure supplement 1) and for analyzing gene sets (e.g. Figure 6 – figure supplement 3D). For global correlations, Spearman’s rank correlation coefficients were calculated. Significance values for pair-wise comparisons of gene expression levels of gene sets (e.g. Figure 6 – figure supplement 3D) were calculated using the Mann-Whitney test. We also assessed differential gene expression, using the SAMMate RPKM values, for each gene and between WT and *H3^H113N^* strains grown in SC media. Student’s t-test was used to calculate p values with the Benjamini-Hochberg procedure to control the false discovery rate at 0.1. Curve fitting for enzymatic kinetic analysis was performed online using “mycurvefit” tool (https://mycurvefit.com),. Predicted values for 0.03 or 0.05 seconds were within 20% of the maximum values detected and used to calculate the initial velocities, which were then transferred to Graphpad Prism version 5.01 and fitted with the Michaelis-Menten non-linear regression function to determine enzymatic parameters.

## Data availability

Gene expression datasets generated by mRNA-sequencing experiments have been deposited in the NCBI GEO database under the primary accession number GSE100034.

## Acknowledgements

We thank Heather Christofk for useful discussions, Mayo Thompson for proofreading, Marco Morselli for assistance in RNA-seq library preparation, Valeria Culotta for the α-SOD1 antibody and helpful guidance in Sod1 assays, and the UCLA Broad Stem Cell Center Sequencing Core. This work was supported in part by a W. M. Keck Foundation Award to S.K.K. and S.S.M. and NIH grants CA178415 to S.K.K., GM074701 to M.F.C., GM42143 to S.S.M., and CA188592 to M.V. O.A.C was supported by the Whitcome and O.A.C. and C.C. by the Dissertation Year Fellowships from UCLA. N.A. was supported by the NCI Ruth L. Kirschstein NRSA for Individual MD/PhD Degree Fellows F30 CA186619 and NIH training grant T32 GM8042. S.Z. was supported by the Amgen Scholars Program.

## Author contributions

Conceptualization, N.A., O.A.C., M.V., Y.X., and S.K.K.; Methodology, N.A., O.A.C., M.V., Y.X., M.F.C., S.S.M., and S.K.K.; Investigation, N.A., O.A.C., M.V., Y.X., C.C., S.S., L.Y., N.M., S.Y., S.Z., J.D., and S.K.K.; Formal Analysis, N.A., O.A.C., and M.V.; Writing – Original Draft, N.A., O.A.C., M.V., and S.K.K.; Writing – Review & Editing, N.A., O.A.C., M.V., Y.X., L.Y., S.S., M.F.C., S.S.M. and S.K.K.; Resources, N.A., O.A.C., M.V., C.C., L.Y., M.F.C., S.S.M., and S.K.K.; Visualization, N.A., O.A.C., and M.V.; Supervision, S.K.K.; Project Administration, S.K.K.; Funding Acquisition, M.F.C., S.S.M., and S.K.K.

## Competing interests

The authors declare no competing interests.

---

**Figure 1 – figure supplement 1.**
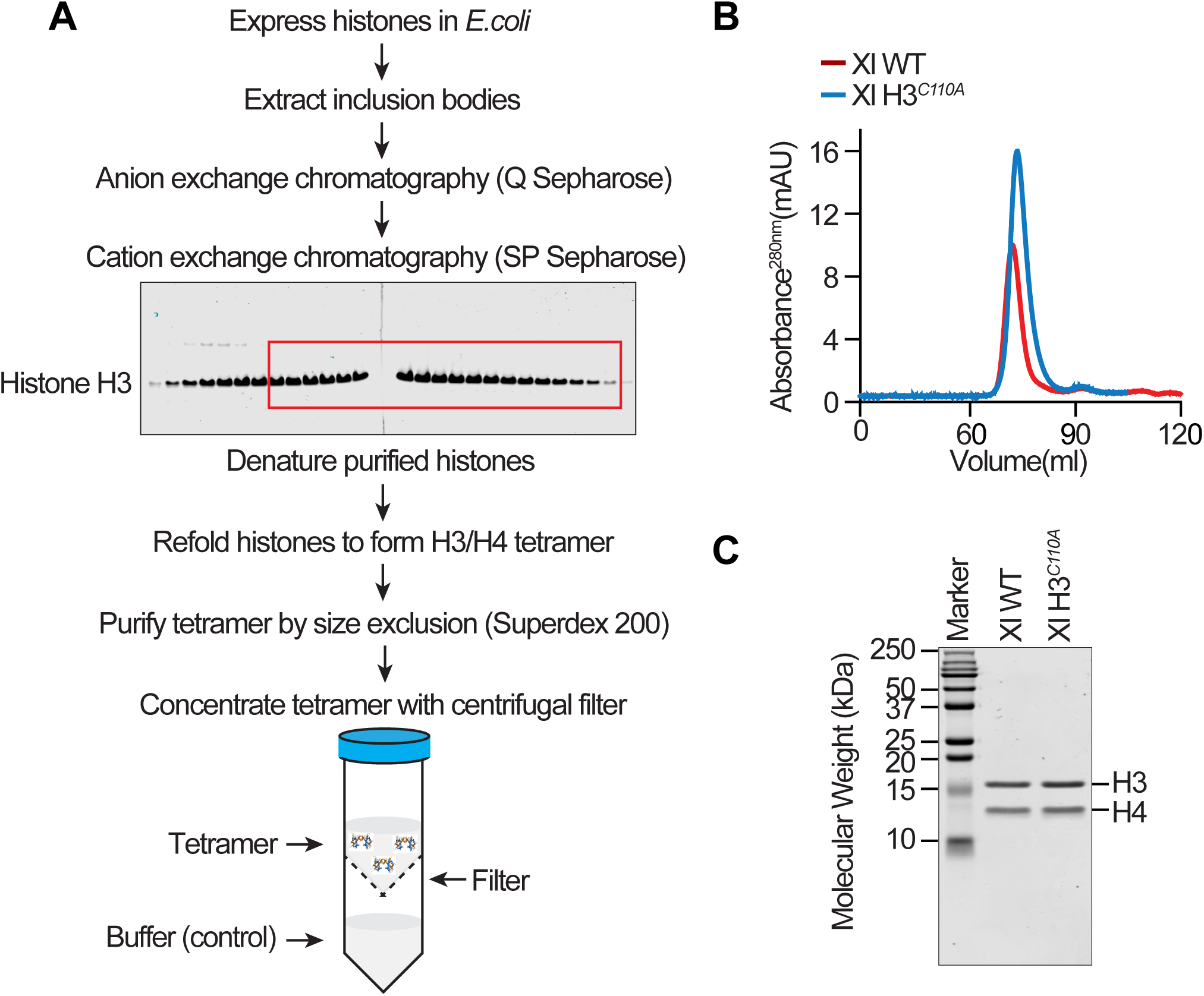
*In vitro* H3-H4 tetramer formation. **(A)** The procedure for preparation of histones and assembly of the *X. laevis* and yeast H3-H4 tetramer is outlined. **(B)** FPLC elution profiles of the indicated *X. laevis* tetramers. **(C)** Image of coomassie-stained PAGE of the indicated purified *X. laevis* tetramers.

**Figure 2 – figure supplement 1.**
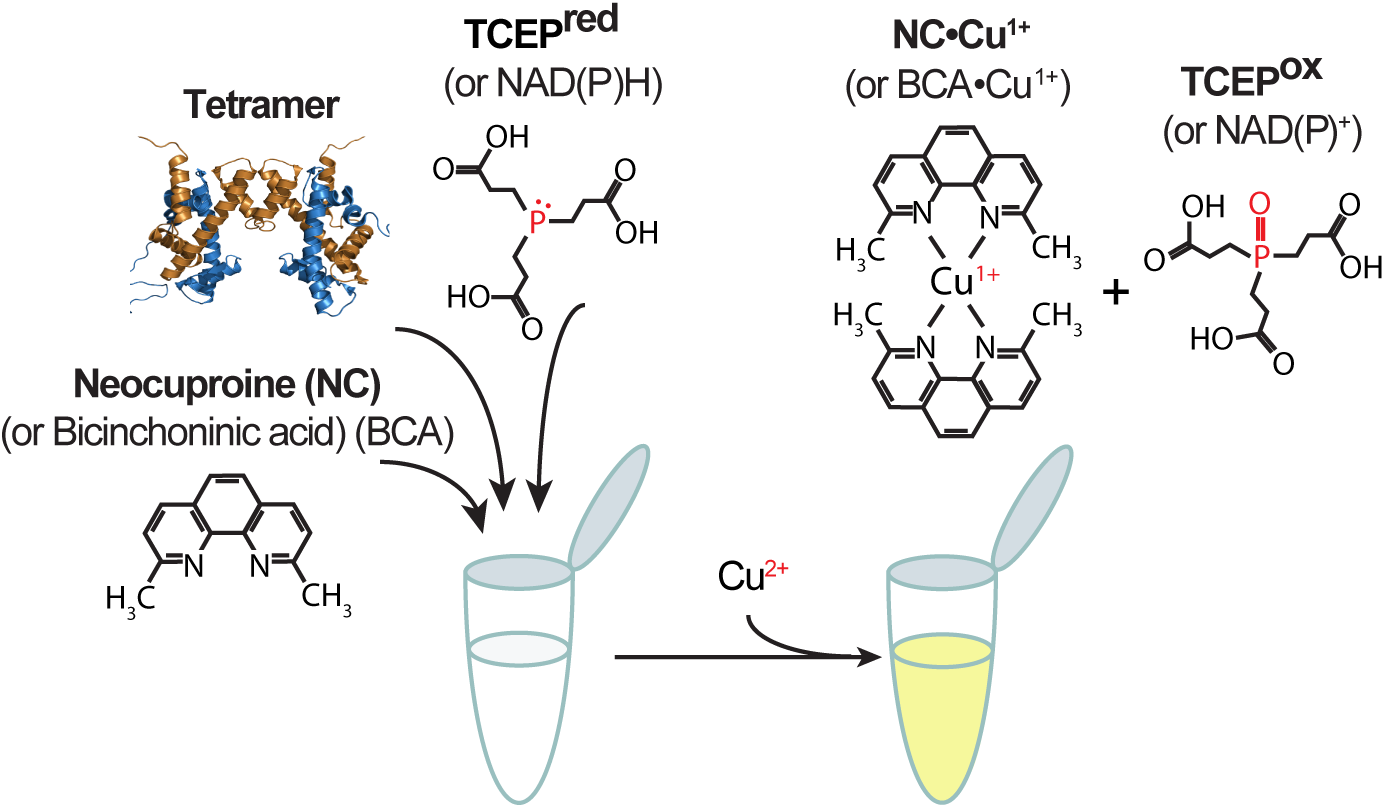
Graphical representation of the copper reductase assay.

**Figure 2 – figure supplement 2.**
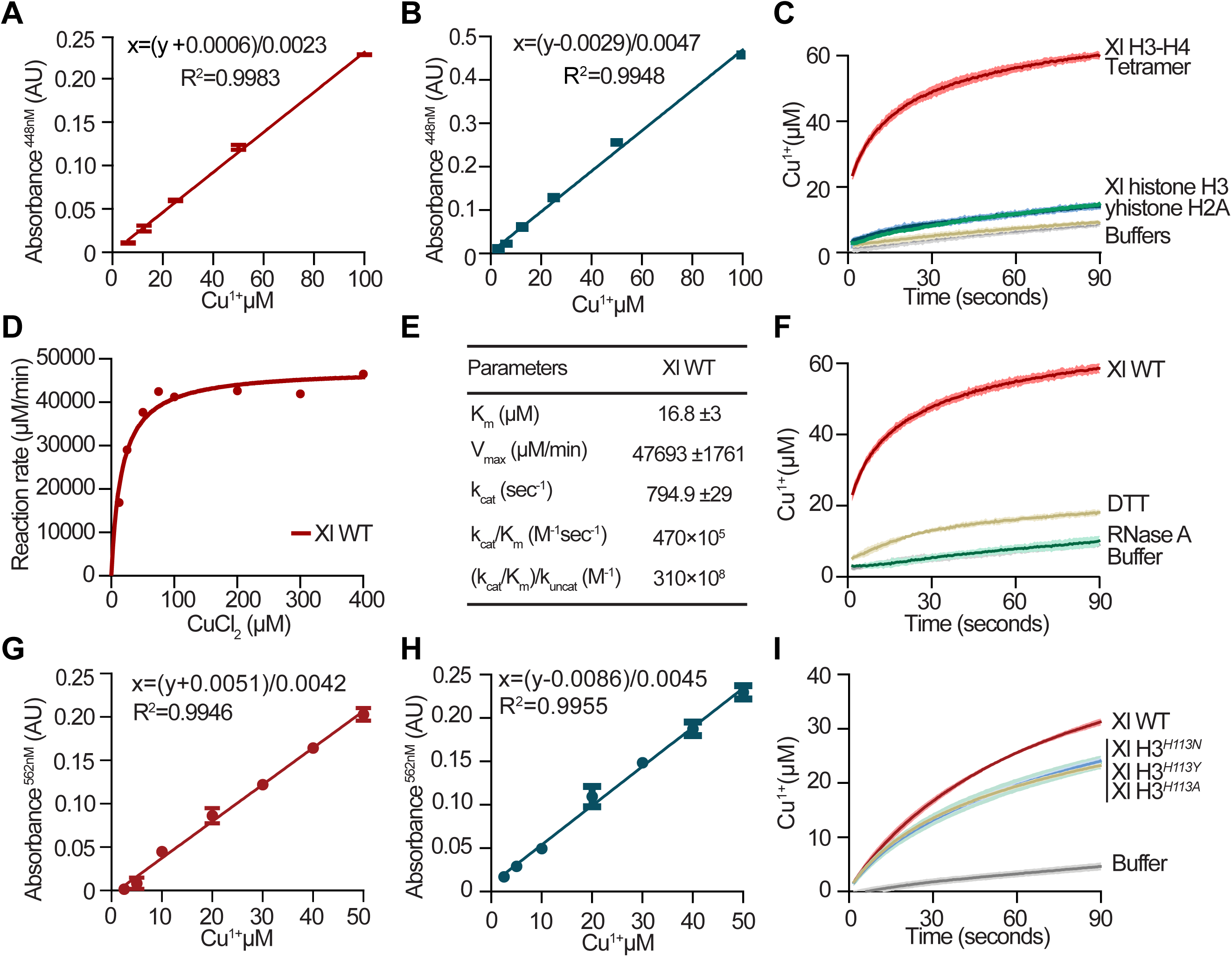
*In vitro* copper reductase assay with the *Xenopus laevis* H3-H4 tetramer. **(A)** Standard curve for Cu^1+^ detection for copper reductase assay conditions used with *X. laevis* tetramers with increasing amounts of CuCl_2_-Tricine pH 7.5 at a ratio 1:10 in presence of 2 mM β-Mercaptoethanol as reducing agent (see Methods). Error bars indicate mean ± SD of three replicate measurements. **(B)** Same as (A) but for reactions with increasing amounts of CuCl_2_-Tricine pH 7.5 at a ratio 1:15 (see Methods). **(C)** Progress curve of copper reduction using 30 µM TCEP as the electron donor with 1 µM of *X. laevis* tetramer, equimolar monomeric *X. laevis* histone H3, and equimolar monomeric *S. cerevisiae* histone H2A. **(D)** Henri-Michaelis-Menten curves of initial velocities of reactions with the *X. laevis* tetramer. **(E)** Calculated enzymatic parameters of the tetramer from the curve in (D). **(F)** Same as (C) but with *X. laevis* tetramer, or equimolar RNase A or DTT. **(G)** Same as (A) but for reactions with increasing amounts of CuCl_2_-ADA pH 8 at a ratio 1:1 (see Methods). **(H)** Same as (A) but for reactions with increasing amounts of CuCl_2_-NTA pH 7.6 at a ratio 1:1 (see Methods). **(I)** Progress curves of copper reduction by 5 µM of the indicated *X. laevis* tetramers in presence of 200 µM TCEP and 0.5 mM CuCl_2_ in 0.5 mM NTA pH 7.6.

**Figure 3 – figure supplement 1.**
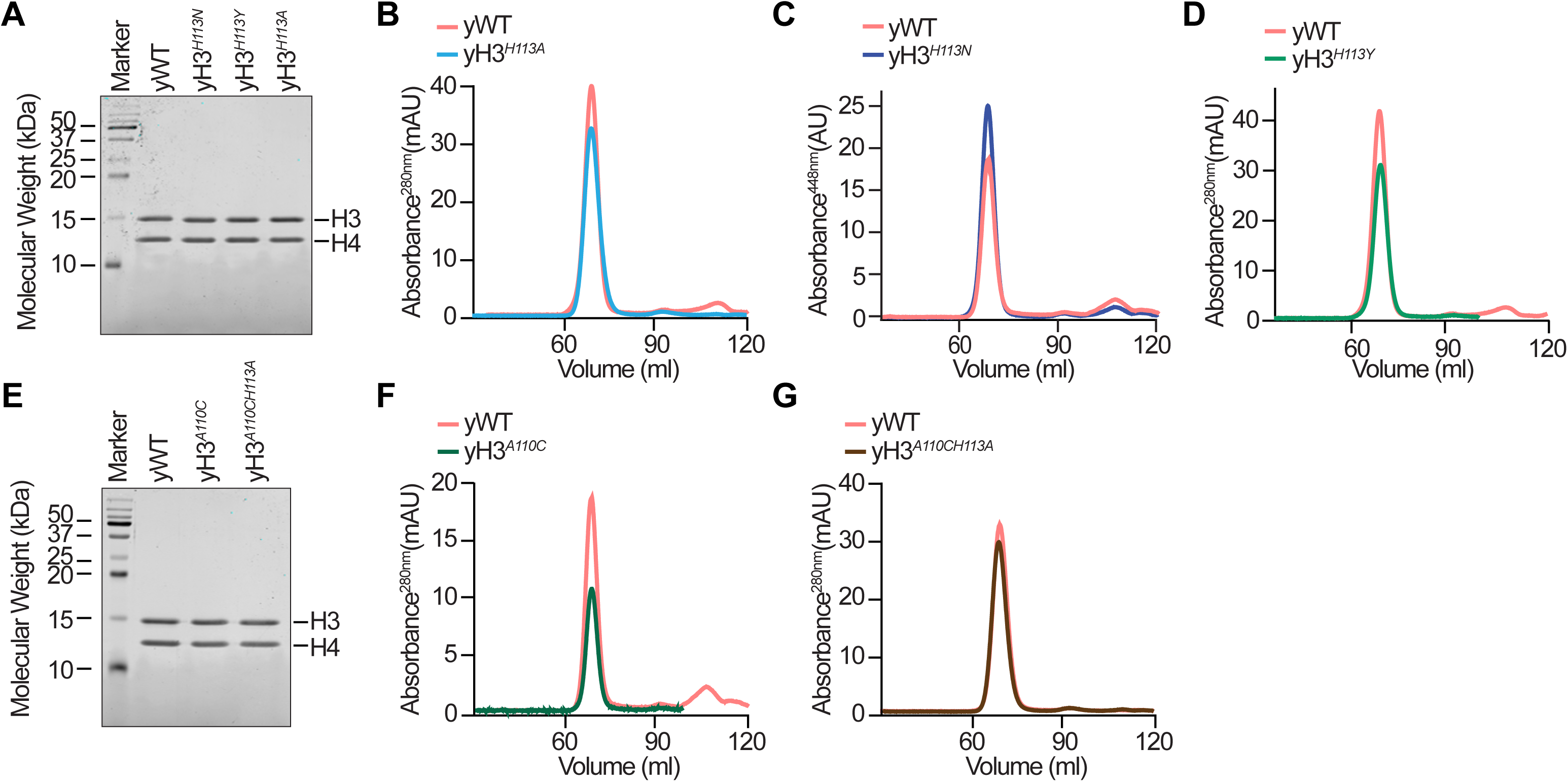
Assembly of *S. cerevisiae* WT and mutant H3-H4 tetramers. **(A)** Image of coomassie-stained PAGE of the indicated purified *S. cerevisiae* tetramers. **(B-D)** FPLC elution profiles of the indicated *S. cerevisiae* tetramers. **(E)** Same as (A). **(F-G)** Same as (B).

**Figure 5—figure supplement 1.**
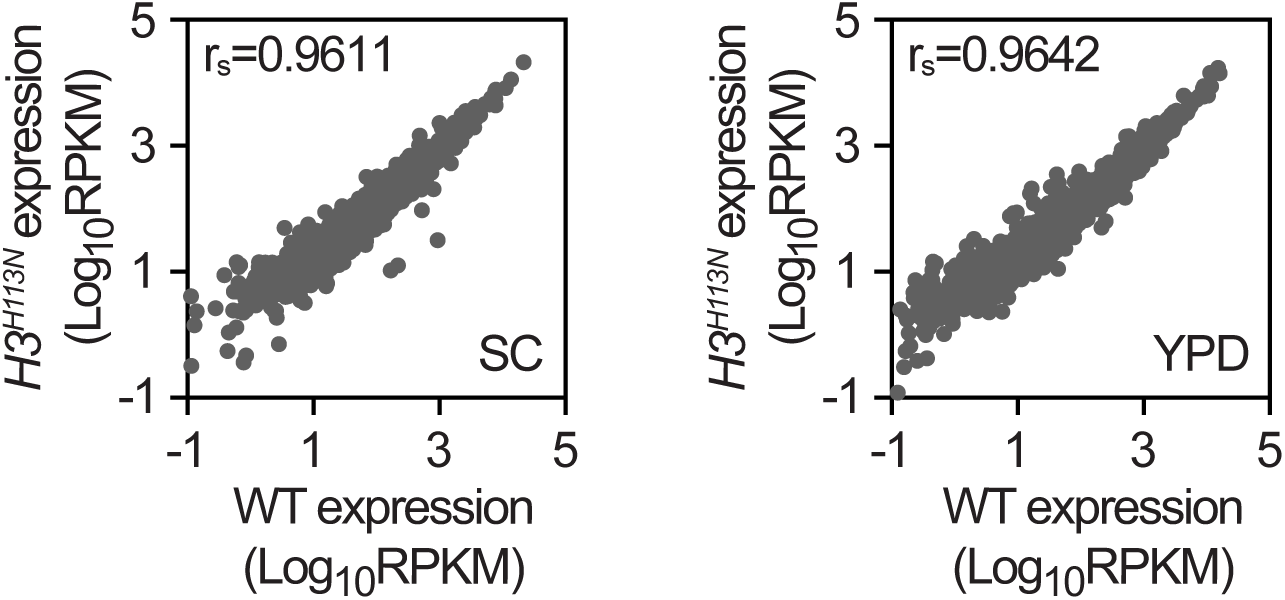
Mutation of H3H113 does not result in global gene expression changes. Scatterplots of average global gene expression values from exponentially growing cells in fermentative media (SC, n=3 and YPD, n=2), with Spearman’s rank correlation coefficients (r_s_) as indicated.

**Figure 6—figure supplement 1.**
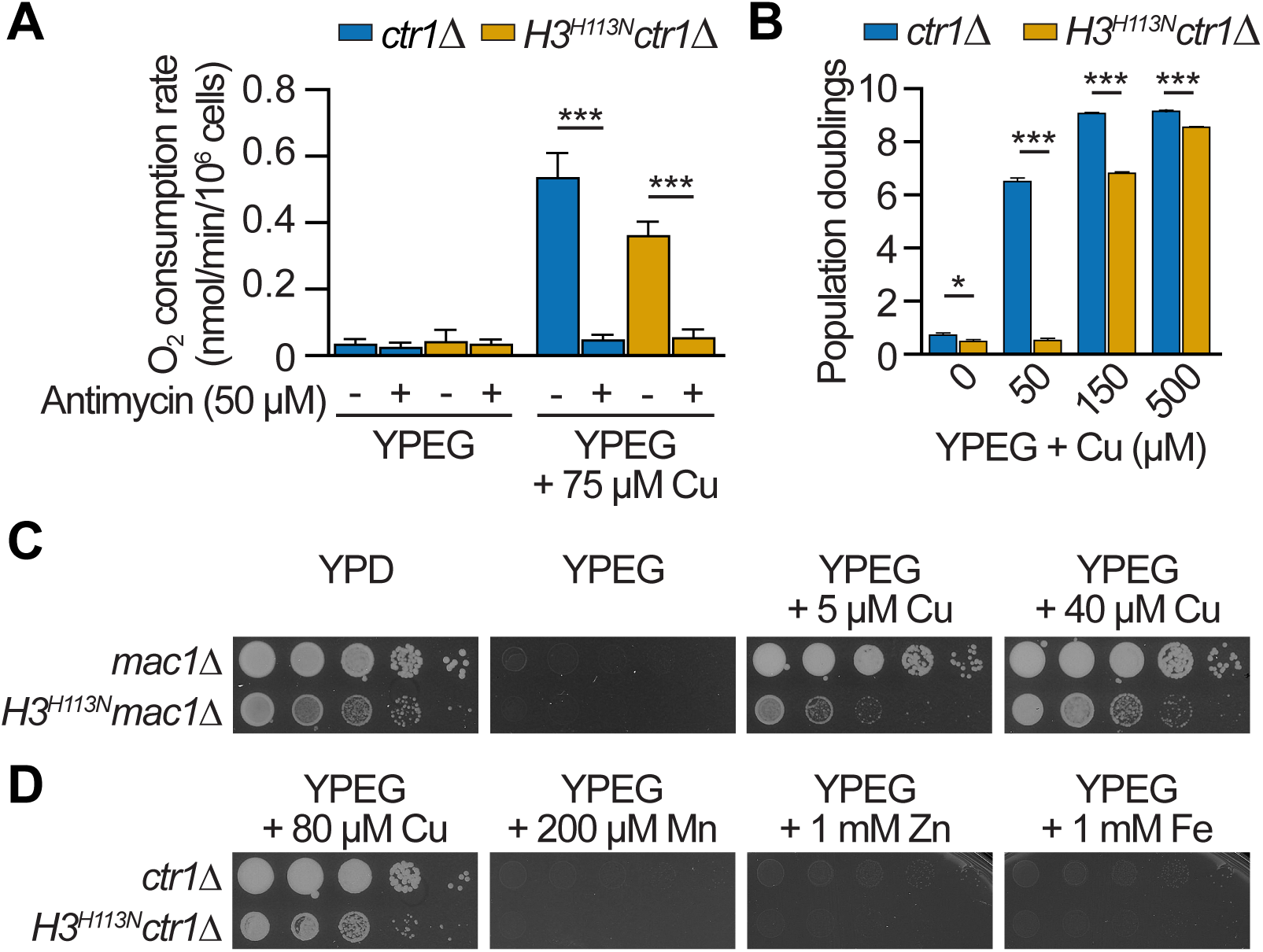
The *H3H113N* mutation diminishes copper utilization for mitochondrial respiration. **(A)** Bar graphs show mean oxygen consumption rates ± SD of the indicated strains from three independent experiments after 6 hrs in liquid YPEG with indicated amount of CuSO_4_ in presence or absence of antimycin A, a mitochondrial complex III inhibitor. Note that cells were pre-loaded with copper prior to growth in YPEG. **(B)** Growth during 48 hrs in liquid YPEG with increasing amounts of CuSO_4_. Bars show mean population doublings ± SD from three independent experiments done concurrently with the same batch of media. **(C)** Spot test assays in fermentative (YPD) or respiratory media (YPEG) with the indicated amounts of CuSO_4_. Baseline copper concentration in YP media is ∼1 µM. **(D)** Same as (C) except with the indicated amounts of MnSO_4_, ZnSO_4_, or FeCl_3_. Note that the *ctr1*Δ strain is respiratory deficient and cannot be rescued by exogenous manganese, zinc or iron. *P≤0.05, ***P≤0.001.

**Figure 6—figure supplement 2.**
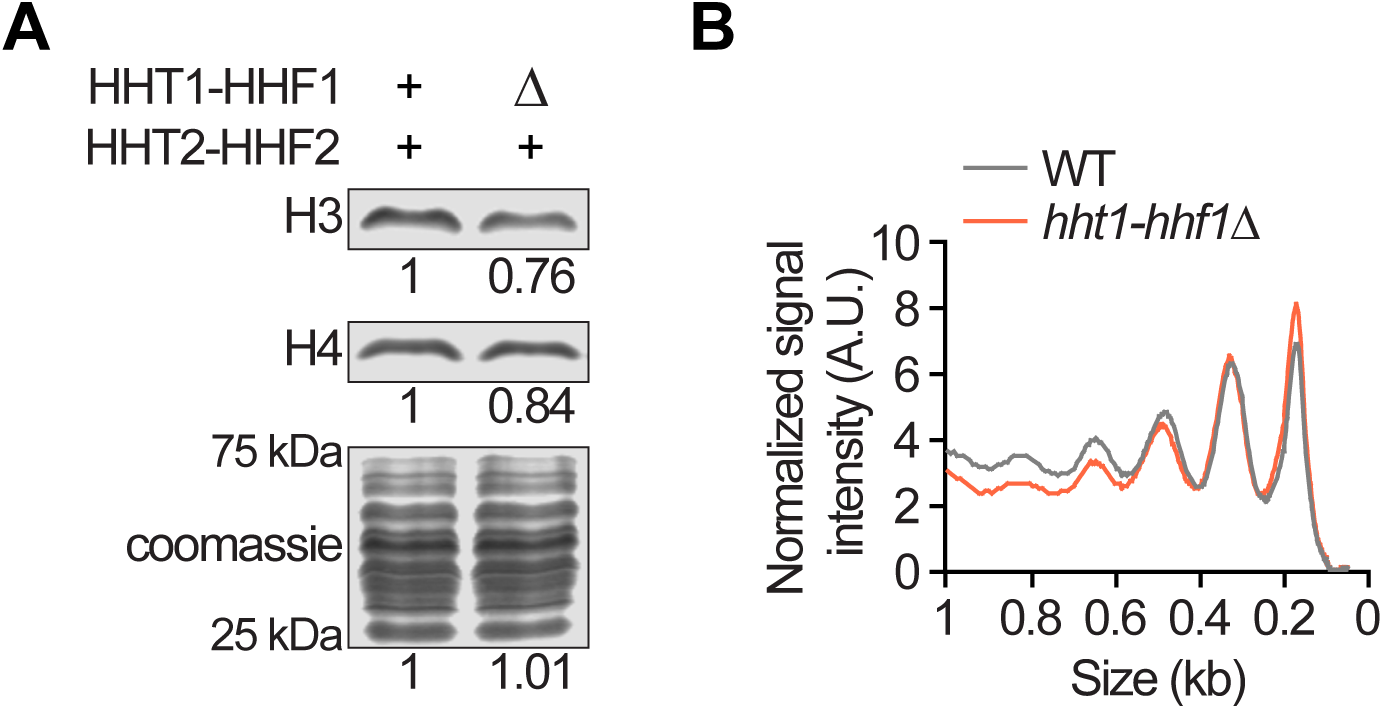
Decrease of the dosage of WT histone H3 and H4 genes. **(A)** Western blots of histone H3 and H4 and coomassie-stained 15% SDS-PAGE, including the relative signal intensity for the indicated strains. **(B)** Signal intensity profiles from chromatin digested with 0.1 units of MNase. A.U.: arbitrary units.

**Figure 6—figure supplement 3.**
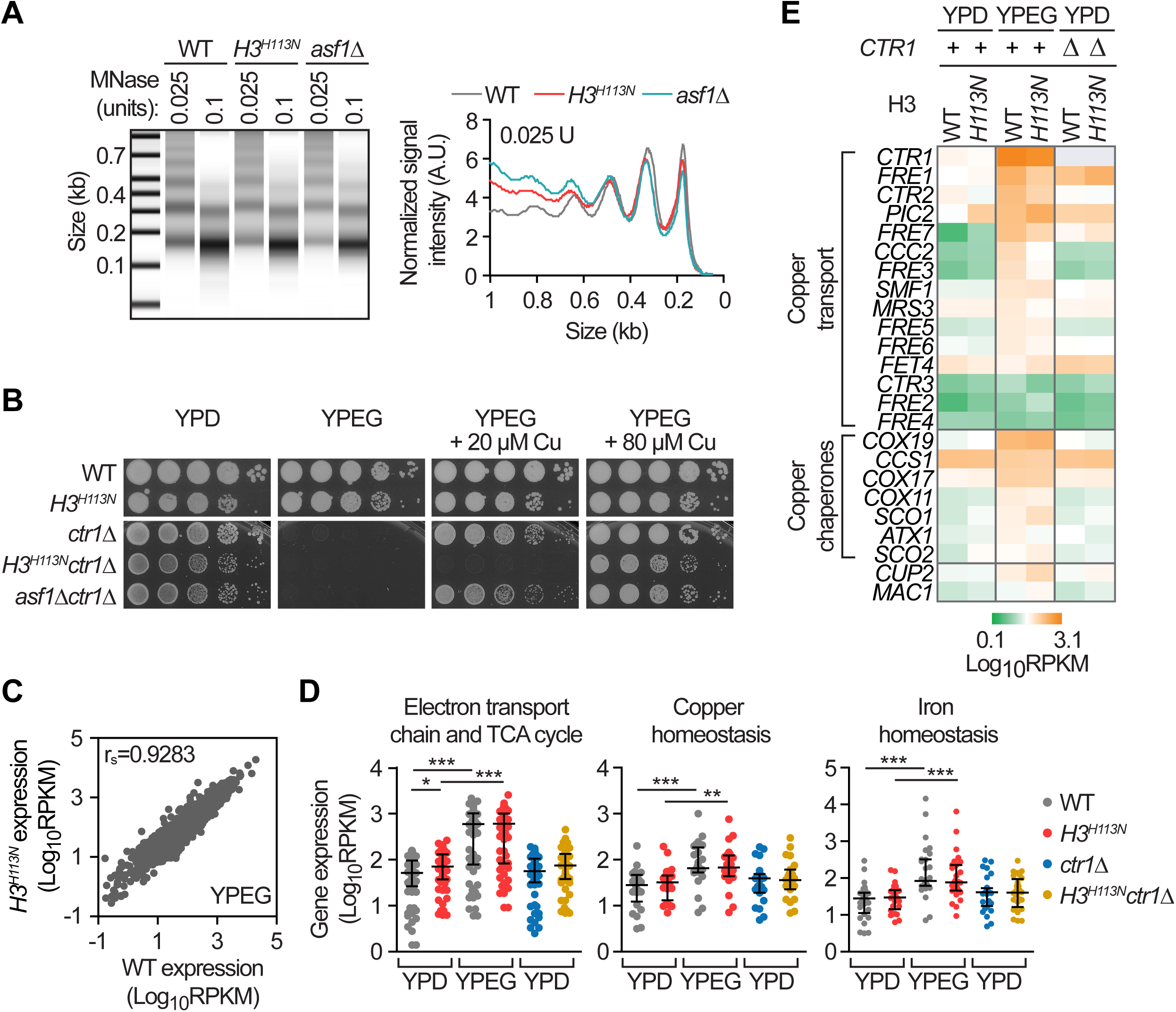
The *H3H113N* mutation does not significantly affect chromatin accessibility or gene expression in respiratory media. **(A)** Quantitative representation of the signal intensity profiles (right) from chromatin digested with the indicated amounts of MNase (left) from exponentially growing cells in SC. A.U.: arbitrary units. **(B)** Spot test assays in fermentative (YPD) or respiratory media (YPEG) with the indicated amounts of CuSO_4_. Baseline copper concentration in YP media is ∼1 µM. **(C)** Scatterplot of average global gene expression values from exponentially growing cells in respiratory media (YPEG, n=2), with Spearman’s rank correlation coefficients (r_s_) as indicated. **(D)** Average mRNA expression levels for three gene sets comparing fermentative (YPD) and respiratory (YPEG) growth conditions from two independent experiments. Each dot is the average RPKM values for an individual gene within the sets and the central lines and whiskers represent the median and interquartile range. **(E)** Heat map of average mRNA expression levels for copper homeostasis genes (data summarized in panel D) from two independent experiments. Genes have been sub-grouped based on known functions of the gene products and ordered vertically within the groups based on the RPKM values of the WT strain in YPEG medium (3^rd^ column). *P≤0.05, ***P≤0.001.

**Figure 6—figure supplement 4.**
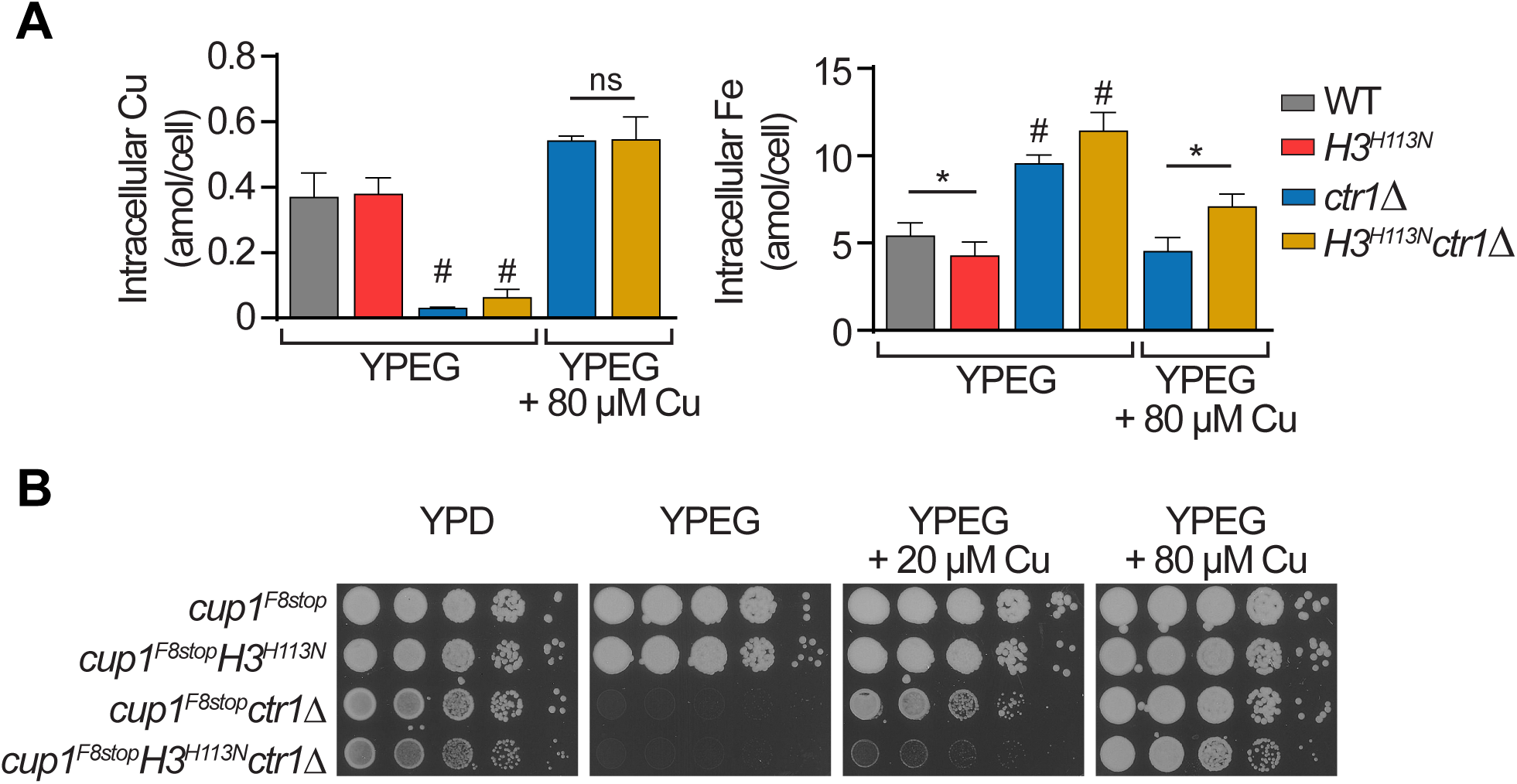
Disruptions in copper utilization are not accounted for by changes in copper content or sequestration by Cup1. **(A)** Intracellular copper (left) and iron (right) content of cells grown in the indicated media for 3-4 doublings for WT and *H3^H113N^* and 12 hrs for *ctr1*Δ strains. Bar graphs represent mean ± SD from 3-6 replicate cultures. ^#^The *ctr1*Δ strains, which do not grow in non-fermentable media, were incubated in YPEG and assessed for metal content for reference. **(B)** Spot test assays in fermentative (YPD) or respiratory media (YPEG) with the indicated amounts of CuSO_4_. Baseline copper concentration in YP media is ∼1 µM. *P≤0.05.

**Figure 6—figure supplement 5.**
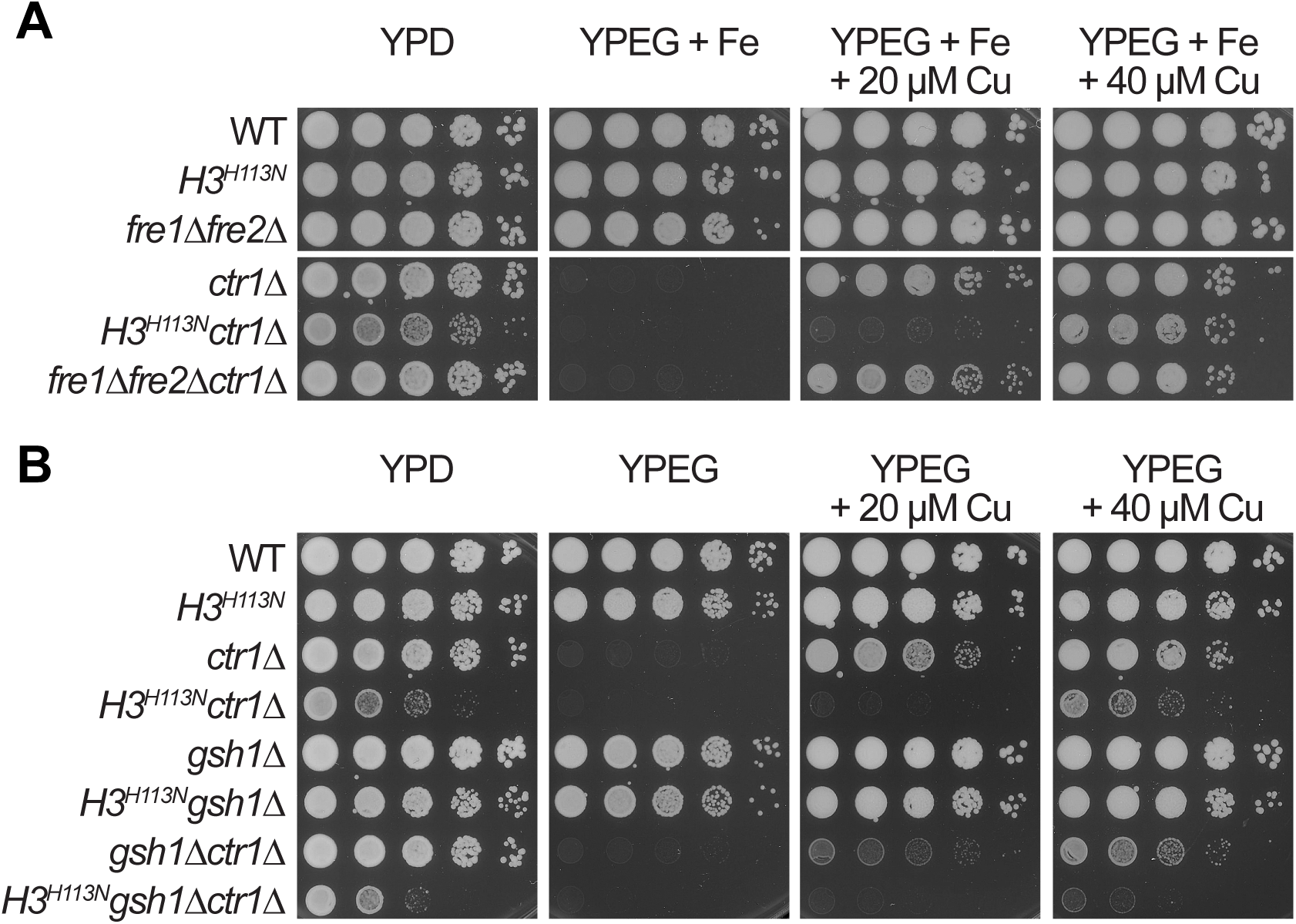
Disruption of cell surface metalloreductases or glutathione synthesis affect copper utilization differently than H3H113 mutations. (A-B) Spot test assays in fermentative (YPD) or respiratory media (YPEG) with the indicated amounts of CuSO_4_. Baseline copper concentration in YP media is ∼1 µM.

**Figure 7—figure supplement 1.**
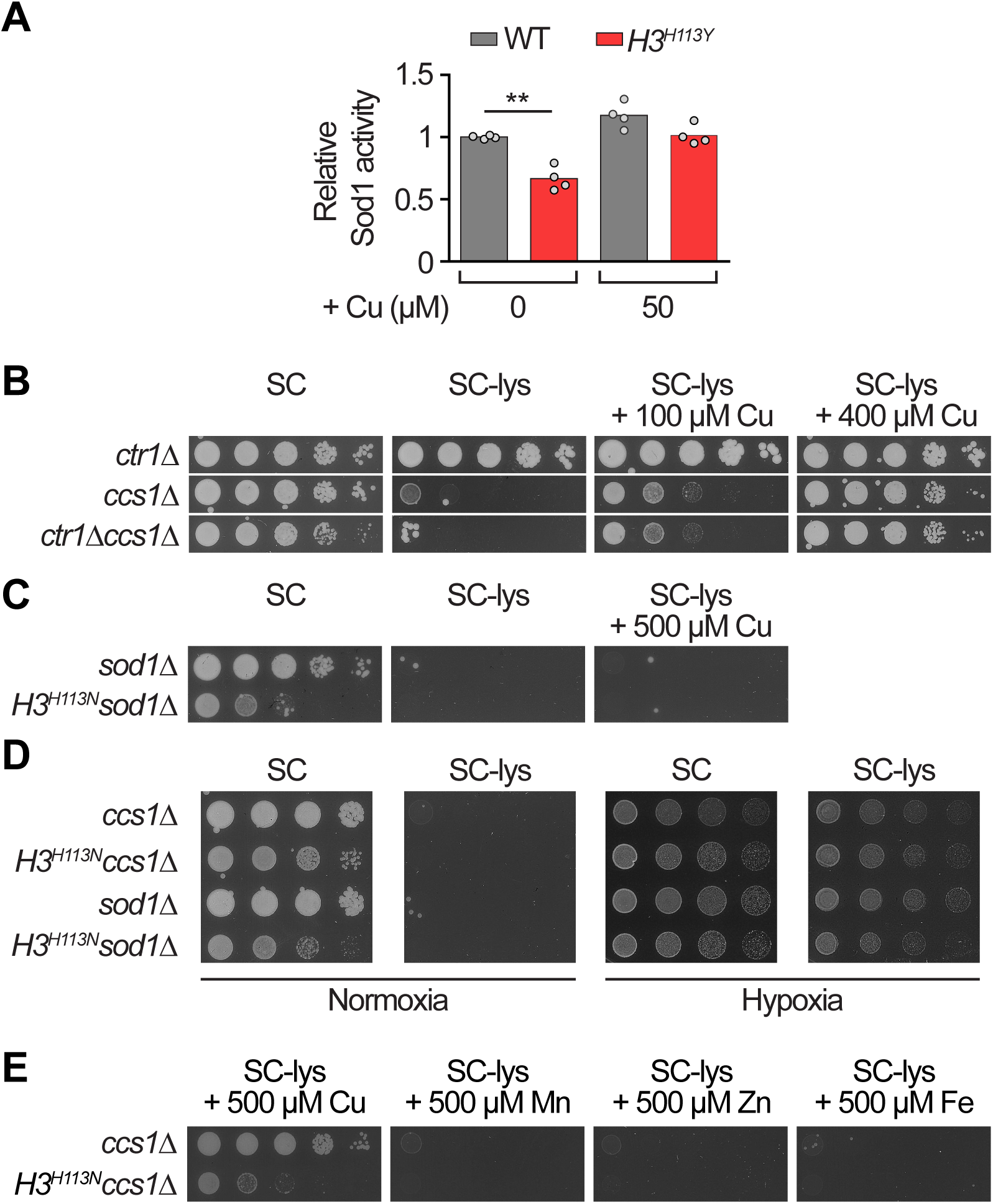
The *H3H113N* mutation diminishes copper utilization for Sod1 activity. **(A)** Normalized relative signal intensities of Sod1 in-gel activity assays for cells grown in SC medium with the indicated amounts of additional CuSO_4_ (representative gel image in Figure 7A). Bars are the sample means and each dot is an independent experiment (n = 4). Baseline copper concentration in SC medium is ∼0.16 µM. **(B-C)** Spot test assays of the indicated strains in fermentative (SC) media with or without lysine and with the indicated amounts of CuSO_4_. **(D)** Same as (B). Hypoxia was generated with an air-tight chamber and an anaerobic gas producing sachet. **(E)** Same as (B) except with the indicated amounts of MnSO_4_, ZnSO_4_, or FeCl_3_. Note that *ccs1*Δ or *sod1*Δ renders cells auxotrophic for lysine and cannot be rescued by exogenous manganese, zinc or iron. **P≤0.01.

**Figure 7—figure supplement 2.**
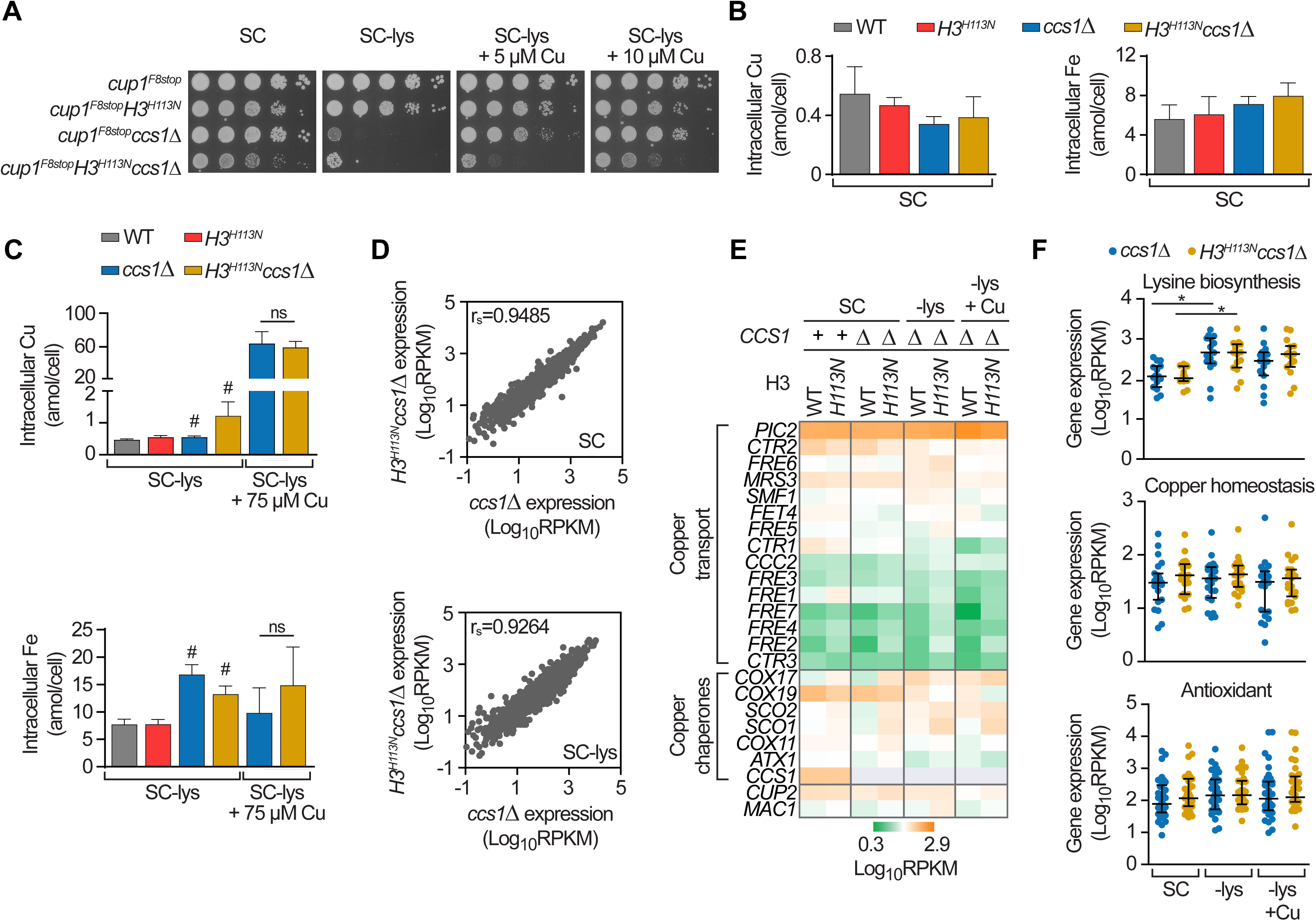
Disruptions in copper dependent Sod1 function are not accounted for by changes in copper content, sequestration by Cup1, or gene expression. **(A)** Spot test assays of the indicated strains in fermentative (SC) media with or without lysine and with the indicated amounts of CuSO_4_. Baseline copper concentration in SC medium is ∼0.16 µM. **(B)** Intracellular copper (left) and iron (right) content measured by ICP-MS for exponentially growing strains in SC. Data are presented as mean ± SD from 3-6 replicate cultures. **(C)** Intracellular copper and iron content of cells grown in the indicated media for 3-4 doublings for WT and *H3^H113N^* and 24 hrs for *ccs1*Δ strains. Bar graphs represent mean ± SD from 3-6 replicate cultures. ^#^The *ccs1*Δ strains, which grow minimally in SC-lys media, were assessed for metal content for reference. **(D)** Scatterplots of average global gene expression values from exponentially growing cells in SC or SC-lys media from two independent experiments, with Spearman’s rank correlation coefficient (r_s_) as indicated. **(E)** Heat map of average mRNA expression levels for copper homeostasis genes (data summarized in panel F) from two independent experiments. Genes have been sub-grouped based on known functions of the gene products and ordered vertically within the groups based on the RPKM values of the *ccs1*Δ strain in SC-lys medium (5^th^ column). **(F)** Average mRNA expression levels for three gene sets in cells growing exponentially in SC, SC-lys, or SC-lys + 75 µM CuSO_4_ media after 24 hrs from two independent experiments (except n=1 for SC-lys+Cu). *P<0.05.

